# Multipartite super-enhancers function in an orientation-dependent manner

**DOI:** 10.1101/2022.07.14.499999

**Authors:** Mira T. Kassouf, Helena S. Francis, Matthew Gosden, Maria C. Suciu, Damien J. Downes, Caroline Harrold, Martin Larke, Marieke Oudelaar, Lucy Cornell, Joseph Blayney, Jelena Telenius, Barbara Xella, Yuki Shen, Nikolaos Sousos, Jacqueline A. Sharpe, Jacqueline Sloane-Stanley, Andrew Smith, Christian Babbs, Jim R. Hughes, Douglas R. Higgs

## Abstract

Transcriptional enhancers regulate gene expression in a developmental-stage and cell-specific manner. They were originally defined as individual regulatory elements that activate expression regardless of distance and orientation to their cognate genes. Genome-wide studies have shown that the mammalian enhancer landscape is much more complex, with different classes of individual enhancers and clusters of enhancer-like elements combining in additive, synergistic and redundant manners, possibly acting as single, integrated regulatory elements. These so-called super-enhancers are largely defined as clusters of enhancer-like elements which recruit particularly high levels of Mediator and often drive high levels of expression of key lineage-specific genes. Here, we analysed 78 erythroid-specific super-enhancers and showed that, as units, they preferentially interact in a directional manner, to drive expression of their cognate genes. Using the well characterised α-globin super-enhancer, we show that inverting this entire structure severely downregulates α-globin expression and activates flanking genes 5’ of the super-enhancer. Our detailed genetic dissection of the α-globin locus clearly attributes the cluster’s functional directionality to its sequence orientation, demonstrating that, unlike regular enhancers, super-enhancers act in an orientation-dependent manner. Together, these findings identify a novel emergent property of super-enhancers and revise current models by which enhancers are thought to contact and activate their cognate genes.

## Introduction

With ever-increasing analysis of the genomes of a wide range of mammalian species, it has become clear that the enhancer landscape is complex. For example, in mouse and human, there are ∼20,000 genes yet >500,000 elements marked with chromatin signatures indicative of enhancer activity^1–3^. However, only a proportion of these elements act as conventional enhancers in stringent assays^4, 5^. This suggests that such signatures may be associated with a broad range of enhancer-like elements with somewhat different roles. Genome-wide analysis has also shown that elements with enhancer signatures may occur in clusters, which have been variously named, most notably as locus control regions (LCRs)^6^ or super-enhancers^7^. Such clusters are defined by the proximity of their constituent elements and the degree to which they recruit the transcriptional regulatory complex termed Mediator and other transcription factors and chromatin modifications found at enhancers. These compound enhancer elements often drive high levels of gene expression and regulate key lineage-specific genes. Important unanswered questions include whether super-enhancers represent a new, functionally distinct class of regulatory elements and display emergent properties beyond that of individual enhancer elements^8^. To date, these questions have been addressed by deleting single elements or combinations of elements from super-enhancers, however, as yet, a full explanation of how the individual elements of super-enhancers combine to activate gene expression is incomplete^3, 9^.

How enhancers find their target promoters is still not fully understood. In recent years, the regulatory space of enhancers and chromatin activity have been defined by genome compartmentalisation^10, 11^. It is now well-established that genomes are organised into self-interacting fractal chromatin domains ranging from tens to thousands of kilobases (kb) in size^12^. Depending on the method of detection and the mechanism associated with their formation, these are referred to, amongst other terms, as Topologically Associating Domains (TADs) and sub-TADs^13–15^. TADs are thought to result from loop extrusion, a model in which the cohesin complex loads on and tracks along the DNA fibre, extruding it in the process into loops until it gets stalled by the CTCF molecules bound to their cognate recognition sites, mostly in convergent orientation^16, 17^; both cohesin and CTCF are enriched at loop anchors and TAD boundaries^13–15^ and their depletion impact on these structures^18–20^. Whether such structures instruct underlying interactions or simply create a permissive environment to allow enhancers and promoters to interact remains under investigation^10, 21^. Other factors have been implicated in enhancer/promoter interaction and specificity including molecular compatibility, linear spacing^22–26^ as well as the presence and distribution of other *cis*-acting regulatory elements (such as CTCF-binding sites) and processes affecting the 3D structure of the TAD (such as transcription and loop extrusion)^10^. Regulatory element sequence orientation, notably that of CTCF consensus sites, seem to impact on TAD structures and the enhancer accessibility to promoters^15, 27–29^.

It is generally agreed that individual enhancers, by definition, act in an orientation-independent manner^30^. However, it is not known if clusters of enhancer-like elements (such as super-enhancers), working as a unit, act in a similar manner. Of interest, in one set of experiments using large, randomly integrated transgenic inserts derived from bacterial artificial chromosomes, it was found that inversion of the β-globin LCR (a well-characterised super-enhancer) reduced expression of the linked β-globin gene cluster^31^. In addition, when the ý-globin LCR was inserted in either orientation within a group of housekeeping genes, although it activated genes both 5′ and 3′, the upregulated genes changed depending on the orientation of the super-enhancer^32^. Finally, when the complex enhancer cluster lying between the *Kcnj2* gene and the *Sox9* gene was inverted, there was a reciprocal change in expression of these two genes^33^. Together, these observations suggest that, whereas single enhancers act in an orientation independent manner, clusters of enhancers may act as a unit with an encoded bias to the direction in which they activate gene expression. However, it remains unclear whether such directionality is a general feature of super-enhancers and what the underlying mechanisms might be.

Here we analysed the direction of interaction and the effects on gene expression of 78 erythroid-specific multipartite super-enhancers in primary mouse erythroid cells. Using the chromosome conformation capture assay termed NG-Capture C^34^, we find that although they may interact with genes located both 5′ (upstream) and 3′ (downstream), super-enhancers predominantly appear to activate erythroid-specific genes located on only one side of the enhancer cluster. To examine the effect of orientation of such clusters on this observed functional directionality, and whether this effect is encoded within the super-enhancer cluster itself, we have used the mouse α-globin locus as an experimental model. We have previously shown that the cluster of enhancers regulating α-globin expression represents one of the most highly ranked super-enhancers in erythroid cells^35, 36^. The five constituent elements of the mouse alpha-globin super-enhancer (R1, R2, R3, Rm and R4) drive high levels of α-globin (*Hba1* and *Hba2*) expression in erythropoiesis (Figure 2B(i)). Deletion of these elements, individually and in informative combinations, demonstrated a hierarchy in the strength of the five constituent enhancers (R1 40%, R2 50% and R4 10%) which appear to act additively on α-globin RNA expression. Of interest, although R3 and Rm have the signature of enhancers and recruit erythroid-specific transcription factors, they appear to have no intrinsic enhancer activity using conventional enhancer assays^35^. With respect to orientation, although the α-globin super-enhancer has a major influence on the expression of the α-globin genes lying 30 kb 3′, it has little activity on the genes within the ∼165kb α-globin TAD lying 12-35Kb upstream of R1. Here, we found that by simply inverting the entire α-globin super-enhancer, with or without surrounding CTCF binding sites or intervening promoters, the predominant interactions from the super-enhancer change direction and while α-globin expression is severely reduced, expression of the genes lying upstream, 5’ of the super-enhancer, are increased. Together, these findings show that, in contrast to individual enhancers, clusters of enhancers, as found in super-enhancers, may interact and influence gene expression in an orientation dependent manner. This functional polarity is promoter agnostic and rather encoded within the cluster itself.

## Results

### Erythroid super-enhancers activate and interact with their cognate promoters in a directional manner

Within the confines of a topologically associating domain (TAD) or sub-TAD, it is thought that the direction of interaction and activity of any enhancer is determined to some extent by the molecular compatibility and distance between enhancers and their cognate promoters^22–26, 37–39^. In addition, enhancer-promoter interactions may be affected by the distribution of other regulatory elements (e.g. CTCF-binding elements) and processes affecting the 3D structure of the TAD or sub-TAD, such as transcription and loop extrusion^10^. Interestingly, the sequence orientation of regulatory elements such as promoters^40^ and more prominently CTCF binding sites^15, 27–29^ has also been implicated in modifying interactions between individual elements and the formation of chromatin domains. However, when tested, it has been shown that individual enhancers act in an orientation independent manner^30^. To investigate whether clusters of enhancer-like elements similarly act in an orientation independent manner within a TAD or sub-TAD, we focused on super-enhancers (SEs). In particular, we studied 95 erythroid-specific super-enhancers previously identified using the ROSE algorithm^7, 35^. Of these, 15 are a single enhancer and 80 are clusters of two or more enhancers; so called multipartite super-enhancers (SEs) (Figure S1A).

To ask how SEs interact with promoters lying upstream (5’) and downstream (3’), we first defined the domain of interaction (sub-TAD) containing each SE using NG Capture-C^34^. We established viewpoints from 157 individual enhancers embedded within 78 of the multipartite SEs, with at least two viewpoints covering 68 of the 78 clusters (Figure S1B): two SEs could not be included for technical reasons. We next identified erythroid-specific interactions between the erythroid SEs and their cognate promoters within each sub-TAD, comparing NG Capture-C data from primary mouse erythroid cells with those from undifferentiated (non-erythroid) mouse embryonic stem cells (mESCs).

Within the 78 sub-TADs, we identified all NG Capture-C *Dpn*II fragments (15,669 fragments) specifically enriched in erythroid cells (DESeq2, q<0.05)^41^. We next asked how these erythroid-enriched fragments are distributed with respect to the erythroid specific SEs within each sub-TAD as illustrated by one of the 78 erythroid-specific SEs studied here (super-enhancer 25: SE25) (Fig. 1A). In erythroid cells, differentially interacting fragments at the SE25 locus were greatly skewed to the downstream (3’) end of the sub-TAD (Figure 1A and B). In total, we found a similarly skewed distribution of erythroid-specific interactions within the sub-TADs of 54 of 78 (69.2%) erythroid-specific SEs. Thirty SEs showed enrichment of interactions upstream (5’), and 24 showed enrichment downstream (3’).

**Figure 1.**
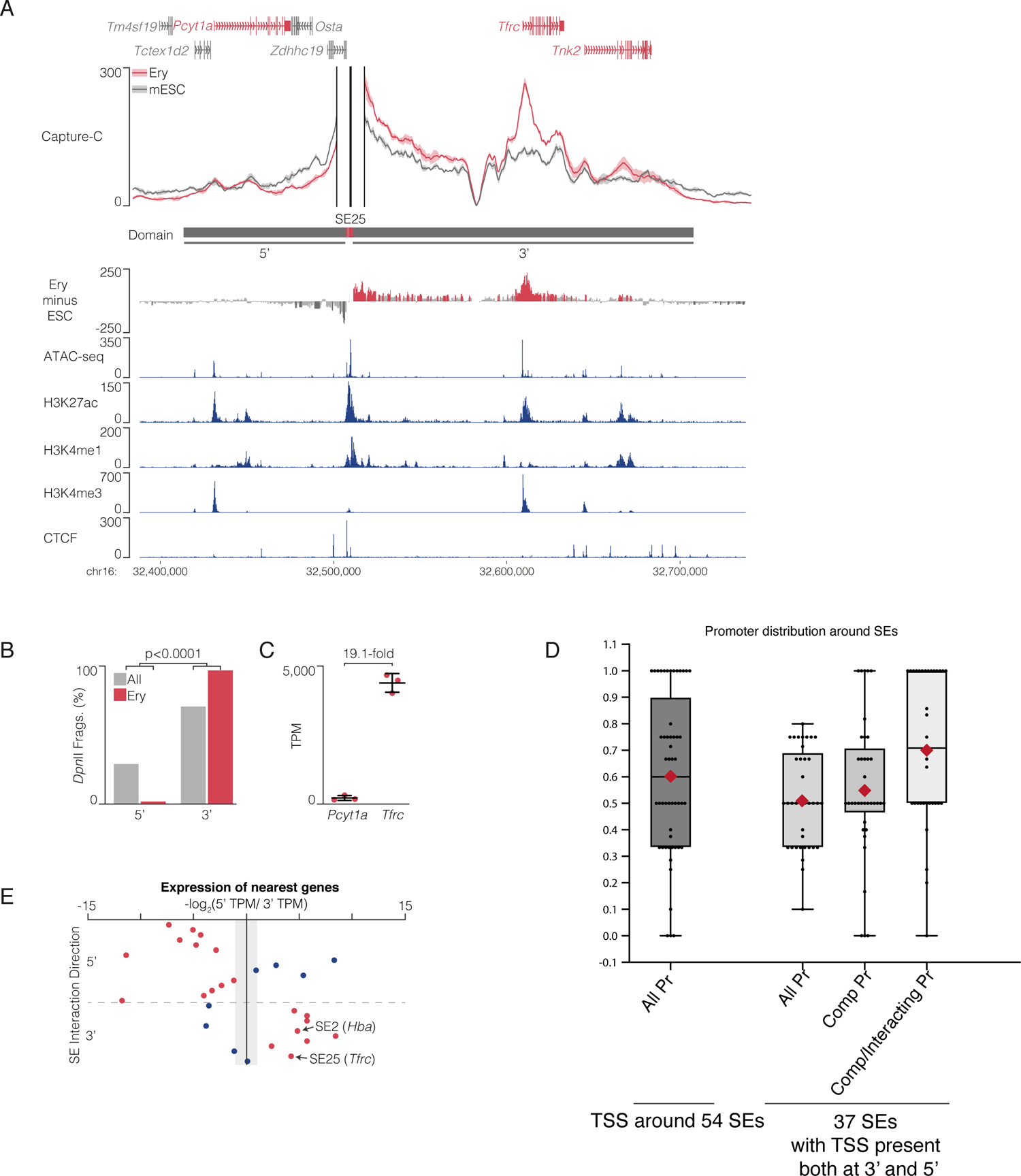
Multipartite super-enhancers show interaction orientation bias linked to active and erythroid-specific genes. (A) NG Capture-C 3C interaction profiles for a representative super-enhancer (SE) (SE25) in mouse ter119^+^ erythroid cells (Ery, red track) and mouse embryonic stem cells (mESC, grey track). Solid lines show means (n=3 independent biological replicates) with one standard deviation (shading) using a 6 kb window. Additional tracks show subtraction (Ery minus ESC) per *Dpn*II fragment, with DESeq2 significantly interacting fragments (p.adj<0.05) represented by dark grey (reduced interactions) or red (increased interactions) peaks. Tracks underneath represent marks of open chromatin (ATAC-seq), and ChIP-seq for active chromatin (H3K27ac), enhancers (H3K4me1), competent promoters (H3K4me3), and CTCF binding. Note, at the top, UCSC gene annotations for genes associated with competent promoters are labelled in red. Region shown is chr16:32,376,000-32,738,000 bp (mm9). (B) Example of the quantitative analysis of interactions showing enrichment in erythroid-specific interactions in SE25 skewed to the 3’ side of the SE: percentage of *Dpn*II fragments (All) and erythroid tissue-specific interacting *Dpn*II fragments (Ery) in the SE25 interaction domain found upstream (5’) or downstream (3’) of the SE. P-value for a Fischer’s exact test with Bonferroni correction. (C) Relative Fold difference in nascent gene expression (TPM, transcripts per million) for the nearest competent 5’ (*Pcyt1a*) and 3’ (*Tfrc*) genes to SE25 shows the higher expressing gene (*Tfrc*) positioned to the 3’ side of the SE. TPM determined by 4sU-seq in maturing terminally differentiating ter119+ erythroid cells (n=3 independent biological samples). (D) The physical distribution of promoters relative to SEs. The distribution is calculated as the ratio of the number of promoters located upstream (5’) or downstream (3’) of the SE (matching the side of the SE interaction skew) over the total number of promoters surrounding that SE. Box plots show maximum and minimum of the data spread (1^st^ and 4^th^ quartiles, whiskers) and the 2^nd^ and 3^rd^ quartiles (boxes). The means of the ratios are indicated by red diamonds. The first box plot (dark grey, All Promoters (All Pr)) represents the distribution of all the promoters (annotated transcription start sites (TSS) based on UCSC) found in all 54 SEs that showed significantly skewed interactions. A mean value of 0.60 indicates that the promoters within this dataset are evenly distributed around SEs. Note that a value above or below 0.5 indicates where the highest number of promoters lie, relative to the SE in question. The following three light grey box plots represent 37 SEs (a subset of the 54 SEs analysed in the first dataset) where promoters (TSS) were found simultaneously upstream (5’) and downstream (3’) of the SE. Analysing this subgroup of SEs avoids any bias arising from SEs with promoters located to one side, only upstream (5’) or downstream of the SE. First light grey box plot (All Promoters, All Pr) represents the annotated promoters surrounding all 37 SEs and shows an even distribution (mean at 0.51) of such promoters upstream (5’) and downstream (3’) of these SEs. The second light grey box plot (Competent Promoters, Comp Pr) accounts for all competent (marked by H3K4me3) promoters with the potential to be activated by the associated SE. This category of promoters is evenly distributed (mean at 0.55) albeit with a shift in the box (representing 50% of the data) above 0.5. The third light grey box plot (Competent/Interacting Pr) shows a skewed distribution of the promoters that are both competent (marked by H3K4me3) and, based on NG Capture-C analysis, interact with SEs in erythroid cells (ratio >0.7), a value above 0.5 indicating a correlation between the skewed pattern of interaction and the distribution of active, erythroid-specific promoters. (E) Relating the direction of SE interactions to gene expression: a representation of SEs (red and blue dots) with skewed direction of interaction towards genes lying upstream (5’) or downstream (3’) and the corresponding relative levels of gene expression of the competent genes lying closest to each of these SEs in the upstream (5’) or downstream (3’) direction. Twenty-eight SEs that are flanked by competent promoters both upstream (5’) and downstream (3’) are ranked based on the skew of their interaction profiles, either to the upstream (5’) (above the horizontal dashed grey line) or the downstream (3’) (below the horizontal dashed grey line). The vertical axis separates relative fold difference in nascent expression (transcripts per million, TPM) of the nearest competent genes to the SEs, calculated as -log2(5’TMP/3’TMP), and allowing the placement of the SE with the highest expression to the 5’ (left of the vertical axis) or 3’ (right of the vertical axis). The grey shaded region on the y-axis indicates less than 2-fold change in expression. Red dots mark the SEs where the interaction direction skew and increase in levels of expression of the nearest genes are concordant. SE25 (*Tfrc* locus), and SE2 (*Hba* locus) are indicated (black arrows).

Skewed interactions could result from a similarly skewed distribution of promoters flanking the SEs within the sub-TADs. To address this, we examined the distribution of all annotated Transcription Start Sites (TSSs) within the 54 sub-TADs that were found to have skewed interactions. We found that all annotated promoters were equally distributed upstream (5’) and downstream (3’) of the erythroid-specific SEs (Figure 1D, All Promoters (All Pr); mean ratio 0.6). To avoid bias, we removed from our analysis any sub-TADs in which SEs are only flanked by promoters on either the upstream (5’) or downstream (3’) sides. In the remaining 37 sub-TADs, where promoters were found both upstream (5’) and downstream (3’) of the SE, we found that they were evenly distributed on either side of the erythroid SEs (Figure 1D, All Promoters (All Pr); mean ratio 0.5). These data show that skewed interactions between SEs and promoters does not result from imbalance in the distribution of promoters around the erythroid SEs.

A bias could also result from the distribution of poised and active (“competent”) promoters compared with silent (non-activatable) promoters. When we analysed the distribution of promoters that are competent (marked by H3K4me3^42^), we found that these are also evenly distributed around the SEs, (Fig 1D and Fig S1C, Comp Pr, mean 0.5). This suggests the 37 SEs analysed could potentially interact equally with accessible promoters located upstream (3’) and/or downstream (5’). Interestingly, we observed a skewed distribution of erythroid-specific NG Capture-C interactions (mean ratio 0.7) only between SEs and their cognate erythroid-specific promoters (Fig 1D, Fig S1C, Comp/Interacting Pr) indicating a potential functional property of these directional interactions.

To investigate this further, we examined whether these directional super-enhancer interactions regulate expression of the genes with which they preferentially interact. Using 4sU-seq generated from terminally differentiating erythroid cells^40^, we compared the nascent expression of the nearest competent genes lying upstream (5’) and downstream (3’) of each super-enhancer. For example, the SE25 locus is flanked by the competent *Pcyt1a* gene upstream (5’) and the competent *Tfrc* gene downstream (3’). The erythroid super-enhancer SE25 (Figure 1A) interacts and activates the *Tfrc* gene but not the equally competent *Pcyt1a* gene (Figure 1C). To examine this using more SEs, we compared expression of the nearest competent upstream (5’) and downstream (3’) gene for the 28 SEs that are flanked by competent promoters on both sides. For 20 of the 28 SEs, the genes with the highest erythroid-specific expression based on the 4sU-seq data correlated with the direction of the SE interaction skew (Fig 1E, red dots). Note how the location of SEs within sub-TADs harbouring the higher expressing genes to the left or the right of the vertical axis matches the 5’ and 3’ directional interaction of their corresponding SEs, above and below the dashed grey line respectively (Fig 1E, red dots). This concordance between higher gene expression and SE interaction skew persists when we narrowed the analysis to 14 SEs that have genes with tissue-specific interactions. When comparing the mean expression, 10 of these SEs show higher expression for genes in the direction of skewed interaction (Fig 1SD). These data show that SEs are preferentially interacting and activating erythroid-specific target genes in a unidirectional manner.

It’s possible that enhancer-promoter compatibility or CTCF position and binding contribute to the observed SE skewing in interaction and gene activation. However, these findings are also consistent with the proposition that clusters of enhancers, that can be promiscuous^43–45^, may regulate gene expression in an orientation-dependent manner.

### Inversion of the major enhancer element within the context of the α-globin regulatory domain has no effect on α-globin expression in a mESC model of erythropoiesis

To understand what underlies the directionality of super-enhancers, we used the well-characterized mouse α-globin cluster as a model (summarized in Figure S2). The α-globin cluster lies within a 165 kb TAD surrounded by widely-expressed genes. When active, in terminally differentiating erythroid cells, the five elements of the α-globin super-enhancer come into close proximity to the α-globin gene promoters in a ∼65 kb sub-TAD flanked by closely apposed, largely convergent CTCF elements^46^. Expression of the α-globin genes, and the widely-expressed flanking genes, can be accurately assessed in mouse models with easily accessible erythroid cells, and in well validated mESC lines that can be differentiated along the erythroid lineage^35, 47, 48^. Here we have generated and analysed a series of informative rearrangements of the mouse α-globin cluster (Figure 2B and see methods) to investigate how the orientation of one of its individual enhancers and the entire super-enhancer interact and activate promoters located upstream or downstream.

Although it is widely accepted that mammalian enhancers act in an orientation independent manner^30^, this has been tested at relatively few complex endogenous loci *in vivo*^3^. To test this within the context of a well-defined regulatory domain, we examined the effect of inverting the strongest constituent enhancer, termed R2, on the formation of the α-globin sub-TAD and gene expression. We generated modified mESC lines and studied the effect of this modification on the α-globin genes in erythroid cells derived from an *in vitro* hematopoietic differentiation system^48^ (Figure 2A, EB-derived Ery). The embryoid body-derived erythroid cells (EB-derived Ery) are primitive-like when comparing their chromatin accessibility and gene expression profiles to primary primitive erythroblasts, obtained from E10.5 peripheral blood, and definitive erythroid cells (ter119^+^ spleen red blood cells)^48^ (Figure 2A). Most importantly, EB-derived red cells express both embryonic (*Hba-x*) and adult (*Hba-a1/2*) α-globin, both under varying degrees of control from the α-globin enhancer cluster, as described in primary primitive and definitive erythroid cells^35, 36^.

We initially created a mESC model in which the R1 enhancer element was deleted from both alleles (Figure 2B ii, DelR1). As reported previously^35^, the pattern of chromatin accessibility at the remaining enhancers appears identical in the presence or absence of R1 in primary erythroid cells. As in engineered mice, removal of R1 in mESCs led to a ∼40-50% downregulation of α-globin mRNA in EB-derived erythroid cells (Figure 3A, B, DelR1). Therefore, we have generated DelR1 EB-derived erythroid cells whereby α-globin expression almost entirely depends on R2 alone, providing a model in which to study the importance of the orientation of the single major enhancer. To examine the effect of R2 orientation in this setting, we compared erythropoiesis, chromatin accessibility and α-globin expression in EB-derived erythroid cells from mESCs in which R1 was deleted and R2 was either in its native orientation (DelR1) or inverted (DelR1-INVR2) on both alleles (Figures 2B ii, iii). Erythropoiesis, chromatin accessibility and α-globin expression (Hba-a1/2) appeared indistinguishable in two independently derived clones of these two mESC lines, showing that the single major enhancer (R2) acts in an orientation-independent manner in the context of the otherwise intact α-globin regulatory domain (Figure 3A, B, DelR1-INVR2). Furthermore, expression of *Hba-x*, *Nprl3* and erythroid control genes (*CD71* and *pb4.2*) was also unchanged in DelR1-INVR2 erythroid cells (Figure 3B). We therefore concluded that the orientation of the major alpha-globin enhancer (R2) has no effect on the functional directionality of the super-enhancer.

**Figure 2.**
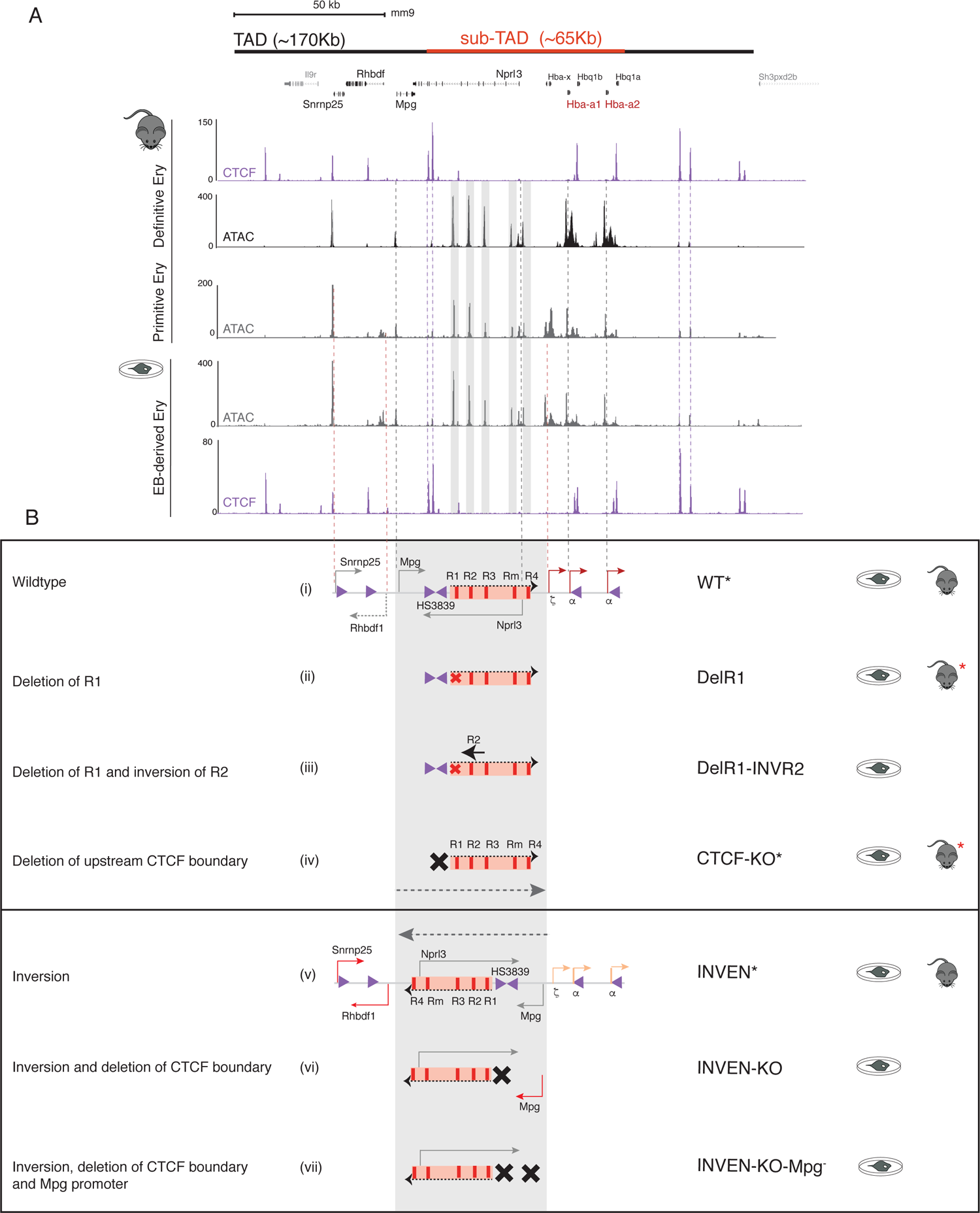
The erythroid systems and mouse models developed and analysed in the study of the inversion and its effects on the α-globin locus. (A) Upper, solid black and red bars represent the α-globin TAD and sub-TAD respectively. RefSeq gene annotation corresponding to the region with the coordinates (mm9) chr11:32,080,000-32,250,000 bp. The lower section shows CTCF occupancy (ChIP-seq) and open chromatin (ATAC-seq) profiles in primary erythroid cells (labelled and marked with the ‘mouse’ schematic) derived from the definitive and primitive lineages as well as EB-derived erythroid cells (labelled and marked with ‘a cell in a dish’ schematic). Note the comparative profiles between the primary primitive and EB-derived erythroid cell profiles, indicative of primitive-like nature of EB-derived erythroid cells. (B) Top box contains schematics representing (i) the wildtype α-globin locus with arrows marking the embryonic (σ) and adult α-globin genes (α) (in red) and flanking genes (*Nprl3, Mpg, Rhbdf1, Snrnp25,* grey arrows) and pointing in the direction of their expression. Orange box represents the SE and the vertical red bars mark the five constituent enhancers (R1-R4). Purple arrows indicate the CTCF-binding sites with the α-globin tested boundary elements labelled (HS38/39). The grey shaded area highlights the region inverted in this study. (ii) schematic of the R1 deletion mutant (red cross, DelR1), (iii) schematic of the R1 deletion (red cross) and R2 inversion (inverted black arrow) mutant DelR1-INVR2, (iv) schematic of the HS3839 boundary deletion mutant (black cross, CTCF-KO). The dashed grey arrow indicates the natural orientation of this piece of DNA with the SE pointing downstream towards the α-globin genes. The lower box contains the inversion models as indicated by the dashed grey arrow pointing upstream away from α-globin genes and towards *Rhbdf1* and *Snrnp25* genes. (v) With the exception of the orientation of the grey-shaded area, and the colour of the arrows representing changes in levels of expression (orange indicating down regulation of the α-globin genes and red arrows indicating upregulation of *Rhbdf1* and *Snrnp25*, the annotation of the elements remains the same as in (i). The dashed black arrow indicates the inverted SE (INVEN). (vi) The INVEN model harbouring the HS3839 CTCF boundary deletion (black cross) and red arrow indicating *Mpg* gene upregulation (INVEN-KO). (viii) The INVEN-KO model harbouring *Mpg* gene knock-out (second black cross, INVEN-KO-Mpg^-^). The black star indicate the models in which *Mpg* gene was also deleted and analysed. Across all models, to the left a summary of what the model represents and to the right, the symbol for the model followed by the schematic indicating the system it was generated in (‘cell in a dish’ for the mESC *in vitro* culture and the ‘mouse’ for *in vivo* mouse model). The red star indicates mouse models previously published.

**Figure 3.**
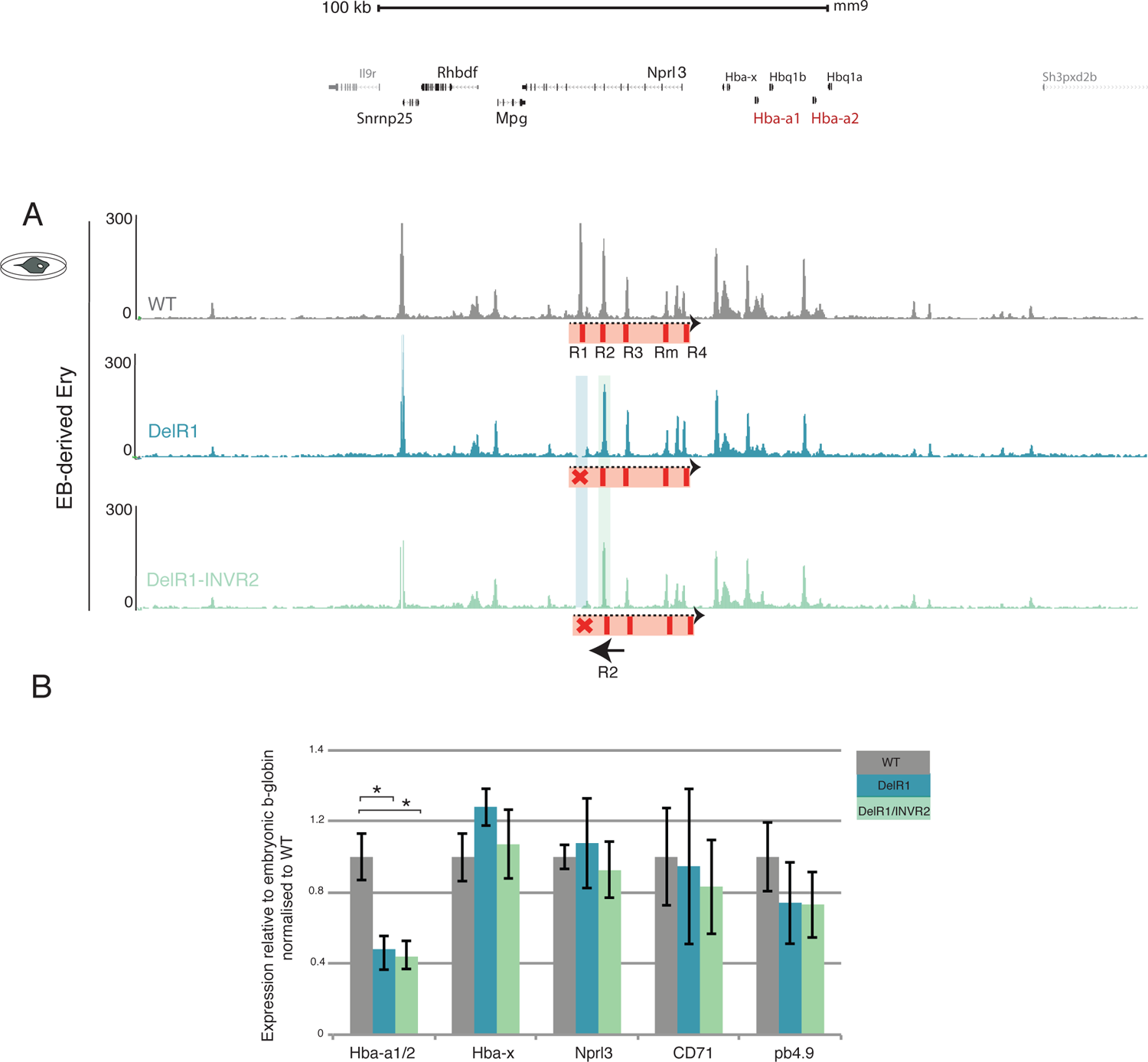
The inversion of the major α-globin enhancer (R2) has no detectable effect on the locus. (A) Upper, RefSeq gene annotation. ATAC-seq tracks show chromatin accessibility profiles in erythroid cells derived from wildtype (WT, grey track) and R1 enhancer deletion (DelR1) mESC models (blue track). Note the absence of the peak corresponding to R1 on the blue track, highlighted with a shaded blue bar. SE schematics as in Figure 2 (i, ii, iii). ATAC-seq track (green) for the DelR1-INVR2 indicates deletion of R1 (shaded blue bar) and intact open chromatin over the inverted enhancer R2 (shaded green bar). (B) Gene expression analysis by real-time qPCR assessing levels of mRNA for *Nprl3, CD71, pb4.2* as controls for the analysed erythroid population, relative to the housekeeping gene 18S and normalised to WT. The α-globin gene expression (*Hba-a1/2 and Hba-x*) was normalised to the embryonic ý-globin gene (*hbb-h1*). At least 4 independent erythroid differentiation experiments were analysed. The bars represent the mean and the error bars indicate the standard deviation (s.d.). *P* values are obtained via an unpaired, two-tailed Student’s t-test. * *P* <0.0001.

### Inversion of the block of all five enhancers within the α-globin regulatory domain down-regulates α-globin expression in a mESC model

The order of the α-globin enhancers within the regulatory domain has been conserved throughout at least 70 million years of evolution^49, 50^. We have shown that the R2 element, at its native position in the genome, exerts equal effect in either orientation on the α-globin expression, in agreement with the established enhancer biology paradigm. However, the question of whether a cluster of linked enhancers within a regulatory domain also acts in a similar orientation-independent manner is not clear. To address this, we asked whether inverting the region encompassing all enhancers (super-enhancer) with respect to the α-globin promoters affects gene expression (Figure 2B i, v, grey shaded area). To do this we inverted a region of the mouse α-globin locus containing all five enhancers within the confines of the highly conserved syntenic regulatory domain (5’-*Rhbdf1*-*Mpg*-*Nprl3*-*Hba-x*-*Hba-a1/2*-*Hba-q1/2*-3’). Since disruption of the *Mpg* or *Nprl3* genes would heighten susceptibility to genotoxicity and lead to developmental abnormalities, respectively^51, 52^, we inverted a 50kb region containing *Mpg*, *Nprl3*, and the α-globin regulatory elements in their entirety. To achieve this, convergent LoxP sites were integrated at flanking insertion sites devoid of chromatin features associated with regulatory elements (Figure S2, shaded orange bars). Inversion of this segment of DNA in mESCs occurred upon expression of Cre recombinase (Figure S2, and Methods). Subsequently, all selectable markers were removed using site-specific recombinases (Methods). The resulting cell line was termed INVEN and the integrity of the genome within and in the flanks of the inversion as well as genome-wide is preserved (Figure S3A). It is important to note that the overall change in distance between the major enhancers (R1 and R2) and the α-globin promoters is negligible (∼5kb) in the inversion. The inversion has no impact on chromatin accessibility over the cluster of enhancers (Figure 4A). CTCF binding across the locus appeared unaltered (Figure 4A) although the interactions of the enhancers change such that 5’ interactions are favoured at the expense of 3’ interactions (Figure 4B). Interestingly, the expression of both embryonic (*Hba-x*) and adult (*Hba-a1/2*) α-globin is reduced to ∼20% of normal in EB-derived INVEN erythroid cells whilst the expression of the surrounding genes (*Rhbdf1* and *Snrnpr25* in their endogenous location and *Mpg* in the repositioned location) is increased (Figure 4C). Despite preserving all the *cis-*elements necessary for full α-globin expression, inverting the super-enhancer significantly downregulates α-globin expression.

**Figure 4.**
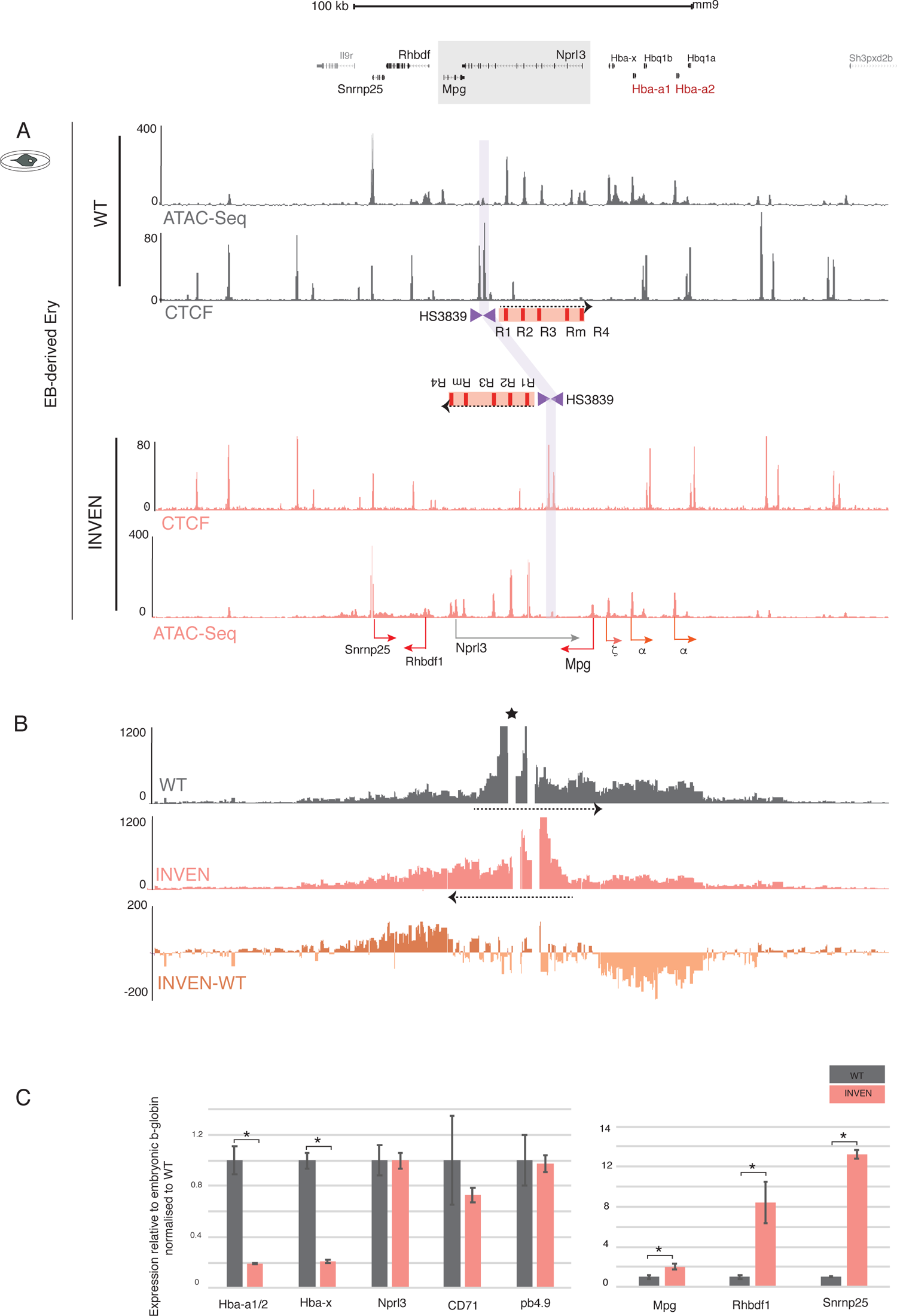
INVEN EB-derived erythroid cells show perturbed SE interaction with the surrounding genes and a change in their expression profile. (A) Upper, RefSeq gene annotation. Lower, normalised (reads per kilobase per million mapped reads, RPKM) and averaged read-densities from 3 independent experiments of ATAC-seq and CTCF ChIP-seq show open chromatin and CTCF occupancy in EB-derived erythroid cells differentiated from WT and INVEN mESCs. For the annotated schematics, refer to Figure 2i and v. The purple shaded bar indicates the position of CTCF boundary element (HS3839) in WT- and INVEN-derived erythroid cells. (B) NG Capture-C interaction profiles in WT-(grey) and INVEN-(pink) derived erythroid cells show normalised and averaged interacting fragment count using a 6 kb window (n=3 independent biological replicates). Additional track shows subtraction (INVEN-WT) per *Dpn*II fragment of significantly interacting fragments using DESeq2 (p.adj<0.05) with light orange for reduced interactions and dark orange for increased interactions in INVEN-derived erythroid cells. The dashed black arrows indicate the direction of the SE in both WT and INVEN models. The star marks the viewpoint (the R1 enhancer) used in the NG Capture-C experiment. (C) Gene expression analysis by real-time qPCR assessing levels of mRNA for red cell makers *Nprl3, CD71, pb4.2*, as well as *Mpg*, *Rhbdf1*, and *Snrnp25*, relative to the housekeeping gene 18S and normalised to WT. The α-globin gene expression (*Hba-a1/2 and Hba-x*) was normalised to the embryonic ý-globin gene (*Hbb-h1*). At least 4 independent erythroid differentiation experiments were analysed. The bars represent the mean and the error bars indicate the standard deviation (s.d.). *P* values are obtained via an unpaired, two-tailed Student’s t-test. * *P* <0.0001.

### Inversion of all five enhancers leads to the phenotype of α−thalassaemia in homozygous mice

To investigate the effect of the inversion on α−globin expression in primary cells, at all stages of development, mESCs heterozygous for the inversion were used to generate a mouse model harbouring the SE inversion described above (Figure 2B v, Figure S2). Mice heterozygous (WT/INVEN) or homozygous (INVEN/INVEN) for the inversion survived to adulthood, bred normally, and were observed at the expected Mendelian ratios (Figure S3B). As in the mESC INVEN model, chromatin accessibility and modifications at the enhancer cluster were unchanged in primary erythroid cells compared to those derived from WT mice (Figure 5A). However, ATAC and H3K4me3 chromatin peaks were reduced at the α-globin promoters (Figure 5A). Mice harbouring the inversion were anaemic, showed a significant reduction in the α/β-globin RNA ratio (Figure 5B), presented with splenomegaly (Figure 5C, Figure S3C) and increased levels of immature red cells (reticulocytes) in their peripheral blood (Figure 5C). The appearance of the blood films and red cell indices (Figure 5C) indicated a typical hypochromic microcytic anaemia associated with α**−**thalassaemia. Flow cytometry of the erythroid populations derived from the bone marrow of homozygous INVEN mice showed no block in erythroid differentiation but a modest expansion of immature populations due to stress erythropoiesis caused by the α−thalassaemia (Figure S3D).

**Figure 5.**
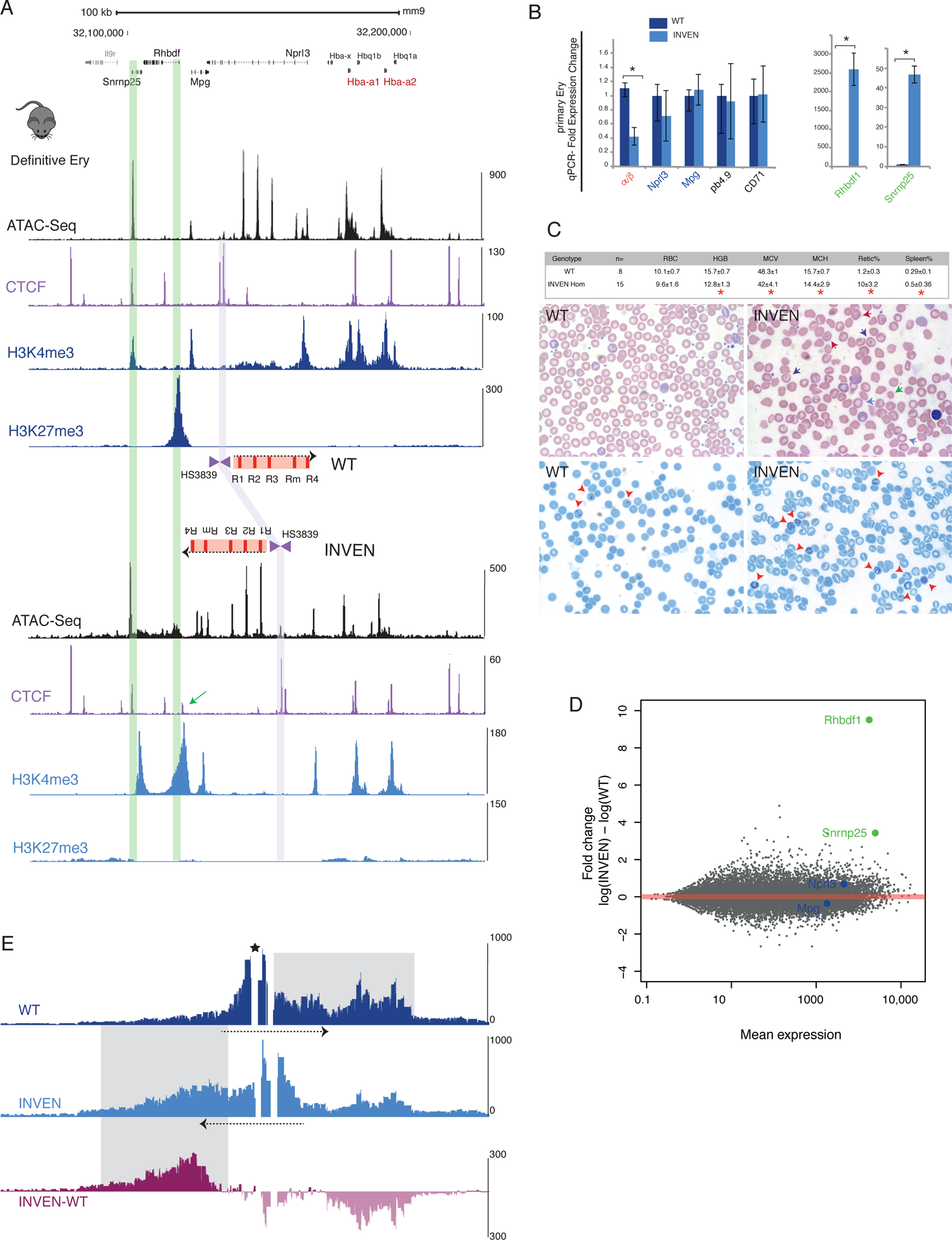
The inversion of the α-globin enhancer cluster (INVEN) perturbs gene expression and chromatin interactions at the α-globin locus and causes an alpha thalassemia phenotype in a mouse model. (A) At the top, scale and RefSeq gene annotation. Normalised (RPKM) and averaged read-densities from 3 independent experiments of ATAC-seq, CTCF, H3K4me3 and H3K27me3 ChIP-seq describe chromatin state in ter119+ spleen-derived definitive erythroid cells from both WT and INVEN mice. For the annotated schematics, refer to Figure 2i and v. The purple shaded bar indicates the position of CTCF boundary element (HS3839) in WT- and INVEN-derived erythroid cells. (B) Gene expression analysis by real-time qPCR assessing levels of mRNA for red cell makers *Nprl3, CD71, pb4.2*, as well as *Mpg*, *Rhbdf1*, and *Snrnp25*, relative to the housekeeping gene 18S and normalised to WT. For the α-globin gene expression, the ratio of α-globin to ý-globin is shown (α/ý). At least 3 independent experiments on mice-derived material were performed. The bars represent the mean and the error bars indicate the standard deviation (s.d.). *P* values are obtained via an unpaired, two-tailed Student’s t-test. * *P* <0.0001. (C) At the top, hematological parameters of red cells: Red Blood Cell (RBC) count, Haemoglobin measurement (HGB), Mean Corpuscular Volume (MCV, fL), Mean Corpuscular Haemoglobin (MCH, g dl^-1^), the reticulocyte percentage (retic%), the spleen weight as a percentage of body weight (Spleen%) are shown for WT and homozygous INEVN mice. *P* values are obtained via an unpaired, two-tailed Student’s t-test. * *P* <0.0001. Below, blood films (upper panels) and Brilliant Cresyl Blue (BCB)-stained blood (lower panels) from WT and homozygous INVEN mice are shown. The INVEN panels show abnormal red blood cells, characteristic of α-thalassemia, indicated in top panel (blood film) by coloured arrows (red: spiky cell membrane (acanthocytes), green: small and round cell (spherocyte), purple: target cells, blue: poorly hemoglobinised hypochromic cells) and lower panel (BCB-stained) with high proportion of immature erythroid cells (reticulocytes, red arrow-heads). (D) MA (log ratio (M) versus average (A)) plot of RNA-seq data derived from WT and INVEN primary erythroid cells. Data represent 3 independent experiments. Mean RNA abundance is plotted on the x-axis and enrichment is plotted on the y axis. Significant upregulation of local genes in the INVEN is highlighted in green (*Rhbdf1*, *Snrnp25*). Blue dots indicate *Nprl3* and *Mpg*, housekeeping genes within the α-globin locus, unaffected by inversion. (E) NG Capture-C interaction profiles in WT (navy) and INVEM (blue) show means (n=3 independent biological replicates) of interacting fragment count using a 6 kb window. Additional track shows subtraction (INVEN-WT) per *Dpn*II fragment of significantly interacting fragments using DESeq2 (p.adj<0.05) with light pink for reduced interactions and dark pink for increased interactions in INVEN-derived erythroid cells. The dashed black arrows indicate the direction of the SE in both WT and INVEN models. The star marks the viewpoint (the R1 enhancer) used in the NG Capture-C experiment.

As the *in vitro* mESC model produces EB-derived primitive-like red cells {HelenaSFrancis:2021fp}, we also harvested E10.5 primitive red cells derived from INVEN mice for comparison. The effect of the inversion in the primary primitive cell context recapitulated the reduced embryonic *Hba-x* and adult α**−**globin (*Hba1/2*) gene expression seen in the *in vitro* model (Figure S4B). E10.5 embryonic erythroid cells show a chromatin accessibility profile similar to that observed in primary INVEN definitive (Figure 5A) and EB-derived INVEN erythroid cells (Figure 4A) marked specifically by a reduced accessibility over the globin promoters and increased accessibility over the *Rhbdf1* and *Snrnp25* genes (Figure S4A). Irrespective of the stage of erythroid development (embryonic or adult), inverting the α**−**globin SE significantly downregulates both *Hba-x* and *Hba1/2* gene expression.

### The inversion reconfigures chromatin conformation in the α−globin regulatory domain and alters the predominant direction of activity of the enhancers

We next asked how inversion of α-globin SE affects chromatin interactions in the α-globin regulatory domain. Using the NG Capture C technique^34^, the self-interacting domain, referred to as a sub-TAD, is normally observed as a ∼65 kb region of increased chromatin interactions, which is formed specifically both in EB-derived and mouse-derived erythroid cells (Figure 4B, 5E, WT tracks). This sub-TAD extends across the entire α−globin cluster with preferential interactions occurring between the α-globin enhancers and promoters ^34, 36, 46, 48^. The erythroid-specific, self-interacting domain is flanked by largely convergent CTCF/cohesin binding sites some of which can act as domain boundaries and restrict the range of interactions of enhancers (Figure S5, CTCFH38 and CTCFH44 viewpoints, WT tracks). The inversion in mESC and mouse models is associated with a newly reconfigured α−globin self-interacting domain whereby decreased interactions between the α−globin enhancers and the α−genes are replaced by newly formed interactions between the α−globin enhancers and the *Rhbdf1* and *Snrnp25* genes (Figure 4B, 5E INVEN-WT track). The newly formed α-globin enhancer interactions were validated by reciprocal capture from the promoters of all genes throughout the 165 kb α-globin TAD including the *Rhbdf1* and *Snrnp25* genes (Figure S6, INVEN-WT track).

In addition to the changes in chromatin accessibility and expression at the α-globin promoters when the α-globin super-enhancer is inverted, we also noted changes in chromatin accessibility, chromatin modifications and expression at *Rhbdf1* and *Snrnp25*. Normally these genes lie upstream (5’) of the α-globin enhancers but in the inversion, they lie downstream of the super-enhancer. Whereas *Rhbdf1* and *Snrnp25* are normally silent or expressed at low levels in WT mice respectively, both are clearly upregulated (2500x and 12x respectively based on normalised RNA-seq read counts) by inverting the SE in definitive erythroid cells derived from the mouse INVEN model (Figure 5B, D). Of particular interest *Rhbdf1* is normally repressed by the Polycomb system in definitive erythroid cells but in the INVEN erythroid cells, the associated chromatin modification (H3K27me3) is completely erased and H3K4me3 acquired as the gene is activated (Figure 5A). We observe similar effects on *Rhbdf1* and *Snrnp25* in embryonic E10.5 and EB-derived primitive erythroid cells (Figure S4B, 4C). Inversion of the super-enhancer did not affect interactions or expression of any other genes lying up to 5.6Mb either side of the 165kb TAD (Figure S7A, B). Thus, inverting the α-globin super-enhancer within the 165 kb TAD redirects enhancer/promoter interactions causing decreased expression of the α−genes, which normally lie downstream, and activation of two genes (*Rhbdf1* and *Snrnp25*) normally lying upstream of the super-enhancer. These findings are consistent with the proposal that the cluster of enhancer-like elements within the super-enhancer works in an orientation-dependent manner within the 165 kb TAD.

### The inversion includes a CTCF insulator element but this does not explain the directionality of the super-enhancer in the α−globin regulatory domain

We previously characterized two CTCF insulator elements (termed HS38/39) which normally lie between the α-globin enhancers and three genes (*Snrnp25*, *Rhbdf1* and *Mpg*) flanking the 3’ edge of the self-interacting domain^47^. To date, these are the only functional insulator elements identified in the α-globin cluster^53^. Removal of this boundary leads to modest activation of the three genes flanking the α-globin sub-TAD but does not affect α-globin expression^47^. This inversion moves HS38/39 from its native position, between the 5’ flanking genes and the SE, such that it now lies between the enhancers, the *Mpg* gene, and the α-globin genes but in the opposite orientation (Figure 2B i, v, Figure S2). Chromatin accessibility and binding of CTCF to this re-positioned element appeared identical to that seen in its native position (Figure 4A, 5A, S4A). Therefore, rather than a change in orientation of the super-enhancer, we asked whether the observed changes in chromatin conformation and gene expression could simply result from changing the position and orientation of this boundary element.

An initial observation argued against this. The engineered rearrangement induced here, which repositioned the insulator element and inverted the super-enhancer, had a much greater effect on interaction and expression of the *Rhbd4* and *Snrnp25* genes than simply deleting the insulator element^47^. This suggests that the predominant effect on these 5’ flanking genes resulted from inverting the super-enhancer rather than deleting the insulator element. To highlight the effect of the inverted super-enhancer without the confounding effect of the boundary elements, we evaluated the role of HS38/39 in the normal locus versus the inverted locus by comparing erythroid cells harbouring HS38/39 deletion in the natural configuration of the locus on both alleles (CTCF-KO/CTCF-KO), to those in which HS38/39 has been removed from both inverted alleles (INVEN-KO/INVEN-KO), in EB-derived erythroid cells resulting from the in vitro differentiation of the engineered models in mESCs (Figure 2B iv, v, vi, Figure 6A). We found that the major changes (chromatin state, expression and interactions) seen in the inversion were not confounded by a boundary effect (comparing data from CTCF-KO to INVEN-KO, Figure 6B, C, S9) nor reversed by removing HS38/39 from the inverted allele (comparing INVEN to INVEN-KO, Figure S8C). In the absence of HS38/39, the direction in which the super-enhancer exerts its effect appears to be encoded intrinsically. In summary, re-positioning of the HS38/39 element is not the cause for reduced interaction between the super-enhancer and the α-globin genes nor it is responsible for the perturbed expression of the surrounding genes, including the upregulation of *Rhbdf1* and *Snrnp25* genes.

**Figure 6.**
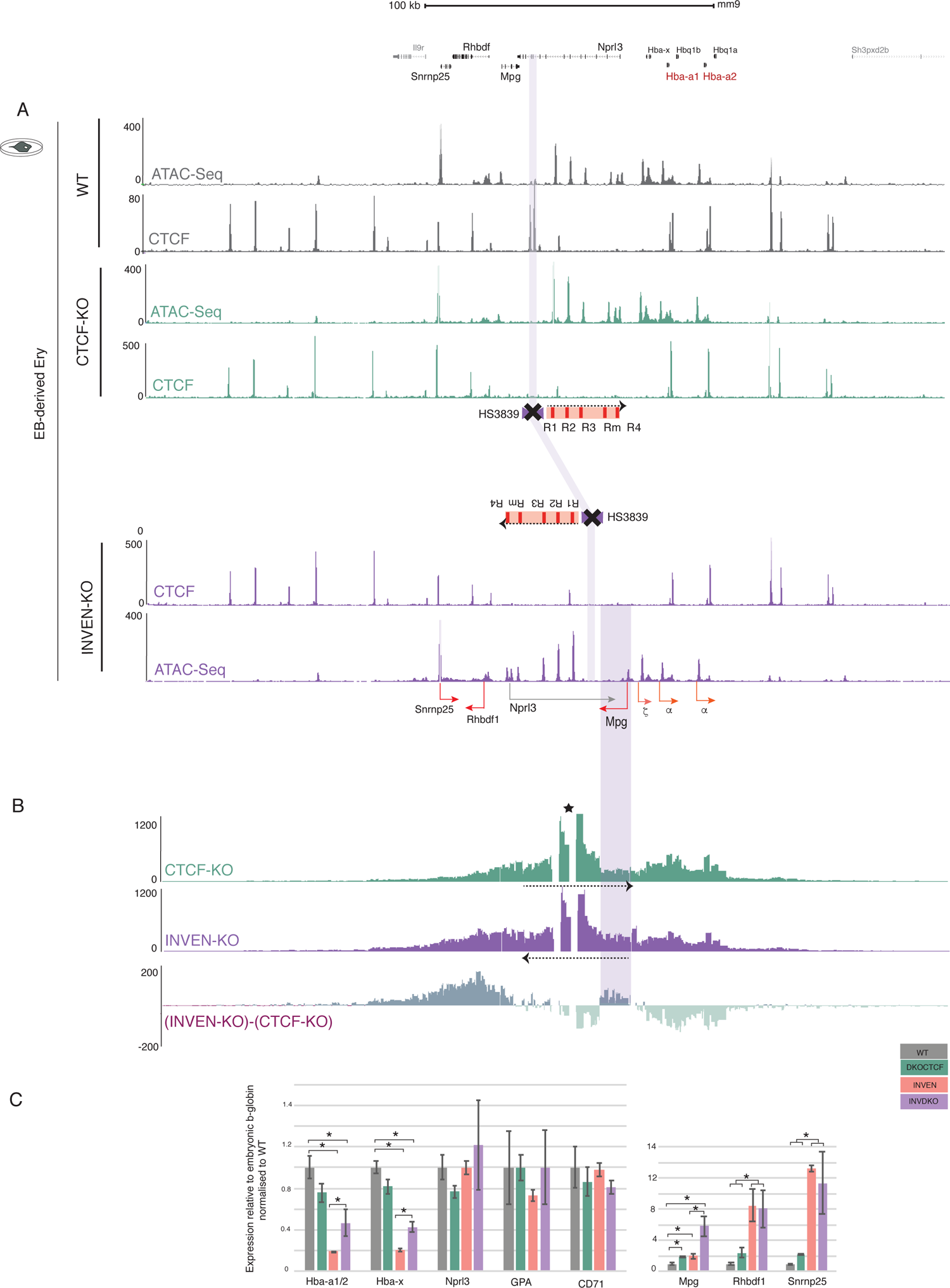
Deletion of the 5’ α-globin CTCF boundary element (HS3839) in INVEN mESC model does not rescue the INVEN phenotype in *in vitro* EB-derived erythroid cells. (A) At the top, scale and RefSeq gene annotation. Normalised (reads per kilobase per million mapped reads, RPKM) and averaged read-densities from 3 independent experiments of ATAC-seq and CTCF ChIP-seq show open chromatin and CTCF occupancy in EB-derived erythroid cells differentiated from WT, CTCF-KO (HS38/39 deleted in WT) and INVEN-KO (HS3839 deleted in INVEN) mESCs. For the annotated schematics, refer to Figure 2i, iv, and vi. The purple shaded bar indicates the position of CTCF boundary element (HS3839) in WT, CTCF-KO and INVEN-KO erythroid cells. Note the absence of the ATAC and ChIP-seq peaks corresponding to CTCF HS3839 in KO models. (B) NG Capture-C 3C interaction profiles in CTCF-KO (green) and INVEN-KO (purple) show means (n=3 independent biological replicates) of interacting fragment count using a 6 kb window. Additional track shows subtraction ((INVEN-KO)-(CTCF-KO)) per *Dpn*II fragment of significantly interacting fragments using DESeq2 (p.adj<0.05) with light green for reduced interactions and dark green for increased interactions in INVEN-KO erythroid cells. The dashed black arrows indicate the direction of the SE in both WT and INVEN models. The star * marks the viewpoint (the R1 enhancer) used in the NG Capture-C experiment. The purple shaded bar indicates *Mpg* gene, located between the SE and the α-globin genes in the INVEN models. Note the increased interactions at the newly positioned *Mpg* in the INVEN-KO. (C) Gene expression analysis by real-time qPCR assessing levels of mRNA for red cell makers *Nprl3, CD71, GPA*, as well as *Mpg*, *Rhbdf1*, and *Snrnp25*, relative to the housekeeping gene 18S and normalised to WT. The α-globin gene expression (*Hba-a1/2 and Hba-x*) was normalised to the embryonic ý-globin gene (*Hbb-h1*). At least 4 independent erythroid differentiation experiments were analysed. The bars represent the mean and the error bars indicate the standard deviation (s.d.). *P* values are obtained via an unpaired, two-tailed Student’s t-test. * *P* <0.0001.

### The repositioned Mpg gene has no impact on enhancer interactions and activity in the inverted locus

A prominent change in interaction we noted was between the super-enhancer and the repositioned *Mpg* promoter in the inverted allele without the boundary element (INVEN-KO) compared with the inversion alone (INVEN) or the WT allele without the boundary element (CTCF-KO), (Figure S8C, Figure 6B). We also reported that this was accompanied by a three-fold increase in expression of *Mpg* (Figure 6C). In both inverted alleles (INVEN and INVEN-KO), the *Mpg* promoter is located between the α-globin super-enhancer and the α-globin genes. We hypothesized that the *Mpg* promoter could act as a new boundary element as proposed for other promoters^13, 54–56^. If so, the *Mpg* promoter may act as a boundary and thereby play a role in reducing α-globin expression and perhaps increasing expression of *Rhbdf1* and *Snrnp25* (Figure 6C). To address this, we deleted the *Mpg* promoter from all the mESC models (WT, CTCF-KO, INVEN, INVEN-KO, Figure 1B i*, iv*, v*, vii) and evaluated the effect of this deletion on expression of the other genes in the landscape (*Rhbdf1*, *Snrnp25*, *Hba-x* and *Hba1/2*) and their interaction profiles. ATAC-seq DNA accessibility profiles confirmed the deletion of the *Mpg* promoter as well as the integrity of all other elements in the domain (Figure 7A). No significant changes in gene expression or interaction profiles were observed when comparing *Mpg*-KO models (Figure 1B i*, iv*, v*, vii) with all the clones they derived from (Figure 7B, C). Despite a moderate increase in α−globin expression, deleting *Mpg* did not restore expression of the α-like globin genes to that observed in the CTCF-KO or WT models (Figure 7C). In summary, the repositioned *Mpg* gene makes little or no contribution to the change in the direction of interaction or activity of the super-enhancer.

**Figure 7.**
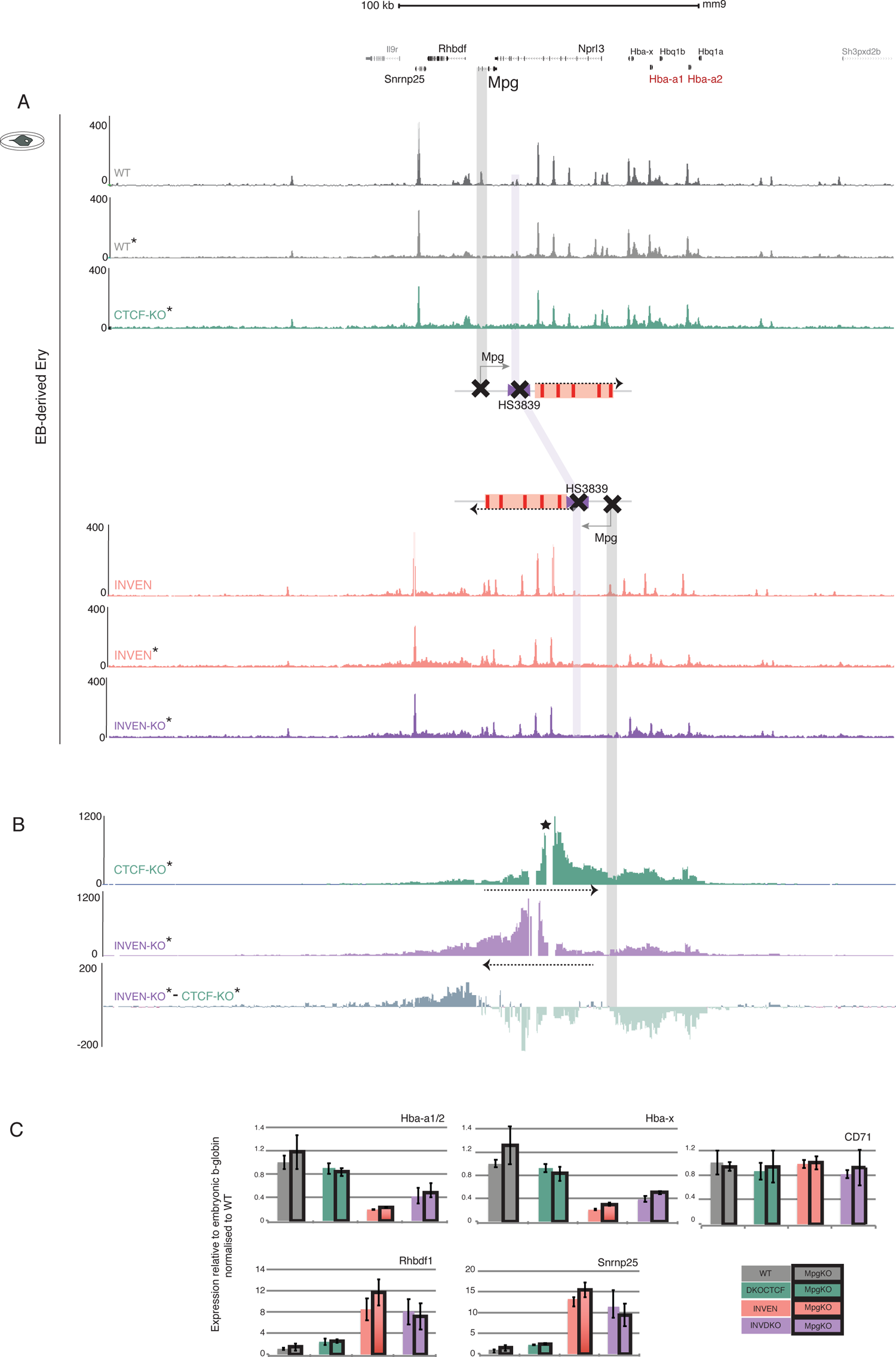
Deletion of the Mpg gene in INVEN mESC models (INVEN and CTCF-KO) does not rescue the INVEN model phenotypes in *in vitro* derived erythroid cells. (A) At the top, scale and RefSeq gene annotation. Normalised (reads per kilobase per million mapped reads, RPKM) and averaged read-densities from 3 independent experiments of ATAC-seq and CTCF ChIP-seq show open chromatin and CTCF occupancy in EB-derived erythroid cells differentiated from WT, WT* (WT with Mpg-KO), CTCF-KO* (CTCF-KO with Mpg-KO) INVEN, INVEN* (INVEN with Mpg-KO) and INVEN-KO* (INVEN-KO with Mpg-KO) mESCs. For the annotated schematics, refer to Figure 2i and v. The purple shaded bar indicates the position of CTCF boundary element (HS3839) in WT- and INVEN-derived erythroid cells. Note the absence of the ATAC peaks corresponding to CTCF HS3839 and *Mpg* gene in KO models (B) NG Capture-C interaction profiles in CTCF-KO* (green) and INVEN-KO* (purple) show means of interacting fragment counts (n=3 independent biological replicates) using a 6 kb window. Additional track shows subtraction ((INVEN-KO)-(CTCF-KO)) per *Dpn*II fragment of significantly interacting fragments using DESeq2 (p.adj<0.05) with light green for reduced interactions and dark green for increased interactions in INVEN-KO erythroid cells. The dashed black arrows indicate the direction of the SE in both WT and INVEN models. The star * marks the viewpoint (the R1 enhancer) used in the NG Capture-C experiment. The purple shaded bar indicates *Mpg* gene, located between the SE and the α-globin genes in the INVEN models. (C) Gene expression analysis by real-time qPCR assessing levels of mRNA for one red cell maker *CD71* as well as *Rhbdf1*, and *Snrnp25*, relative to the housekeeping gene 18S and normalised to WT. The α-globin gene expression (*Hba-a1/2 and Hba-x*) was normalised to the embryonic ý-globin gene (*Hbb-h1*). At least 4 independent erythroid differentiation experiments were analysed. The bars represent the mean and the error bars indicate the standard deviation (s.d.).

## Discussion

There is abundant evidence showing that individual enhancers act in an orientation independent manner in transient transfection assays and when randomly integrated into the genome^57^. Fewer studies have investigated this paradigm by specifically inverting single enhancers at their endogenous locus in their natural genomic context; however, those that have, supported the paradigm of enhancer orientation-independence^58^. Here we have similarly shown that in a model in which the major α-globin enhancer (R2) is inverted in the presence of R3, Rm and R4, there is no detectable change in α-globin RNA expression or in any of the genes within the α-globin sub-TAD. By contrast, to date, the effect of inverting clusters of enhancers (including LCRs and super-enhancers) that might work together to regulate gene expression has not been rigorously tested although some preliminary observations suggest such compound elements may act in an orientation dependent manner^31–33^.

By examining a group of 78 multipartite erythroid super-enhancers, we have shown that although they may interact with, and appear to activate, transcription of genes lying both 5’ and 3’, SEs predominantly interact with promoters distributed in one direction rather than the other. The majority of interactions (in 71% of the SEs studied) between erythroid super-enhancers and genes that are specifically activated and upregulated during erythroid differentiation occur in either one or the other direction from the SE. It is unclear whether this directionality arises from the enhancer-promoter sequence and biochemical affinity^59, 60^, the constraints created by boundary elements and chromatin structures such as TADs^10, 61^ or, in fact, results from information encoded within the super-enhancer ^62^.

Here, detailed analysis of one of the highest scoring erythroid super-enhancers, which regulates **α**-globin expression in erythroid cells^35^, has shown that when considered as a single integrated element, the SE functions in an orientation dependent manner. Importantly, in this case, we were able to rule out any effects of other elements within the α-globin sub-TAD, including CTCF insulator elements and intervening promoters, that might otherwise have confounded this analysis. Together, this suggests that the directionality of the super-enhancer is intrinsic. It is of interest that other observations in the mouse and the orthologous human α-globin cluster also point to the concept that polarity plays some role in the regulation of the cluster. For example, the genes are arranged along the chromosome in the order in which they are expressed in development^46, 63^. This co-linearity is also seen in other multigene clusters most notably in the Hox genes where this phenomenon was first described^64, 65^. In addition, in a wide variety of species, the α-globin genes are duplicated producing alleles with 2, 3 or 4 almost identical tandem copies^50, 66^. In all cases tested, the gene lying closest to the SE is expressed at higher levels than the adjacent gene, which is in turn expressed at higher levels than the next copy and so on^67–70^. A natural mutation which creates a new promoter lying between the super-enhancer and the α-globin genes downregulates α-globin expression, whereas placing this promoter upstream of the super-enhancer, has little effect on α-globin expression^71^. Together these observations suggest that the orientation dependence of the super-enhancer might, at least in part, involve a directional tracking mechanism by which the enhancers and promoters interact.

Current models of enhancer-promoter interactions propose that clusters of enhancers may work together to create a non-membrane bound subnuclear structure containing high concentration of cell specific transcription factors, co-factors including the Mediator complex of transcriptional regulators, and general transcription factors including RNA polymerase II^72–75^. Such structures have been variously labelled as transcription factories, transcriptional hubs and condensates. It is further proposed that promoters contained within such structures are transcribed very efficiently thereby greatly increasing gene expression. The main two activators, R1 and R2, of the α-globin super-enhancer normally lie 30 kb and 26 kb upstream of the duplicated α-globin genes and when inverted they lie 30 kb and 34 kb upstream of these genes. This minor change in distance is very unlikely to have a significant effect on α-gene expression since in the orthologous human cluster a relatively common ∼10kb insertion between the enhancers and the α-genes has no effect on their expression^76^.Consequently, if the super-enhancer simply forms a transcriptional hub, this should be equally accessible to its cognate α-genes in either orientation. This begs the question of why the interaction between the super-enhancer and its cognate genes is orientation dependent.

It has been suggested that transcription could provide a functional link between enhancers and promoters. In the case of the α-globin cluster, short bi-directional eRNAs do not extend between the enhancer and promoter^77^. Furthermore, long non-coding RNAs originating at the elements of the super-enhancer (originally called meRNAs) are normally directed away from the α-globin promoters^77^. It has also been proposed that the interaction between enhancers and promoters is facilitated by loop extrusion mediated by cohesin^16, 17, 28, 78^. Cohesin is indeed enriched at SEs and we have previously observed peaks of cohesin at each element of the mouse α-globin SE, not stalled by CTCF but associated with the Nipbl protein complex which plays a role in loading cohesin^47^. It seems possible that as proposed elsewhere, cohesin may be loaded at super-enhancers from where it could promote loop extrusion and, in some cases, this could occur preferentially in one direction rather than the other^79^. It is of interest that we have shown that the elements in the α-globin SE are not equivalent: two strong conventional enhancers (R1 and R2) are followed by three elements (R3, Rm, R4) with the chromatin signature of enhancers but little or no enhancer activity^35^. This raises the question of whether these elements might provide directionality or timing to the SE, relaying the enhancer activity rather than providing conventional enhancer activity, akin to the previously described “tethering” (GAGA-binding) elements in drosophila^80, 81^. In fact, we have recently reported a detailed functional dissection of the α-globin SE constituent elements where we describe ‘facilitators’ (R3, Rm, R4), a novel form of regulatory elements. We show that ‘facilitators’ are crucial for the full function of the conventional enhancer elements (R1 and R2) within the cluster and most interestingly, that their function is position-rather than sequence-dependent^82^.

In summary, these findings suggest that clusters of enhancer-like elements may act together as an integrated, multipartite element (LCRs and super-enhancers) containing conventional enhancer elements and other elements whose role(s) are not yet clear. In this way, super-enhancers may have emergent properties not present in single conventional enhancers. Importantly, here we show that an unexpected emergent property of a cluster of enhancer-like elements is to act in an orientation dependent manner.

## Acknowledgement

This work was supported by the Medical Research Council (MRC Core Funding number, MR/T014067/1), H.S.F. Wellcome Trust Studentship (109097/Z/15/Z), J.B. Wellcome Trust Studentship (219979/Z/19/Z), L.C. Wellcome Trust Studentship (222843/Z/21/Z).

## Authorship contributions

Methodology, M.T.K., A.S., J.R.H.; Investigation, H.S.F., M.G., M.C.S., C.H., M.L.; Formal Analysis, H.S.F., M.G., M.C.S., C.H., M.L., M.O., L.C., J.B., D.J.D., B.X. N.S.; Software, J.T.; Resources, J.A.S., J.S.S.; Visualisation, M.T.K., H.S.F., D.J.D, M.O., L.C., J.B., Y.S.; Writing-Original Draft, M.T.K., D.R.H.; Writing-Review & Editing, C.B., M.O., D.J.D, H.S.F., J.B.; Supervision, J.R.H.; Conceptualization, M.T.K., D.R.H.; Funding Acquisition, D.R.H.

## Supplementary Figure legends

**Figure S1.**
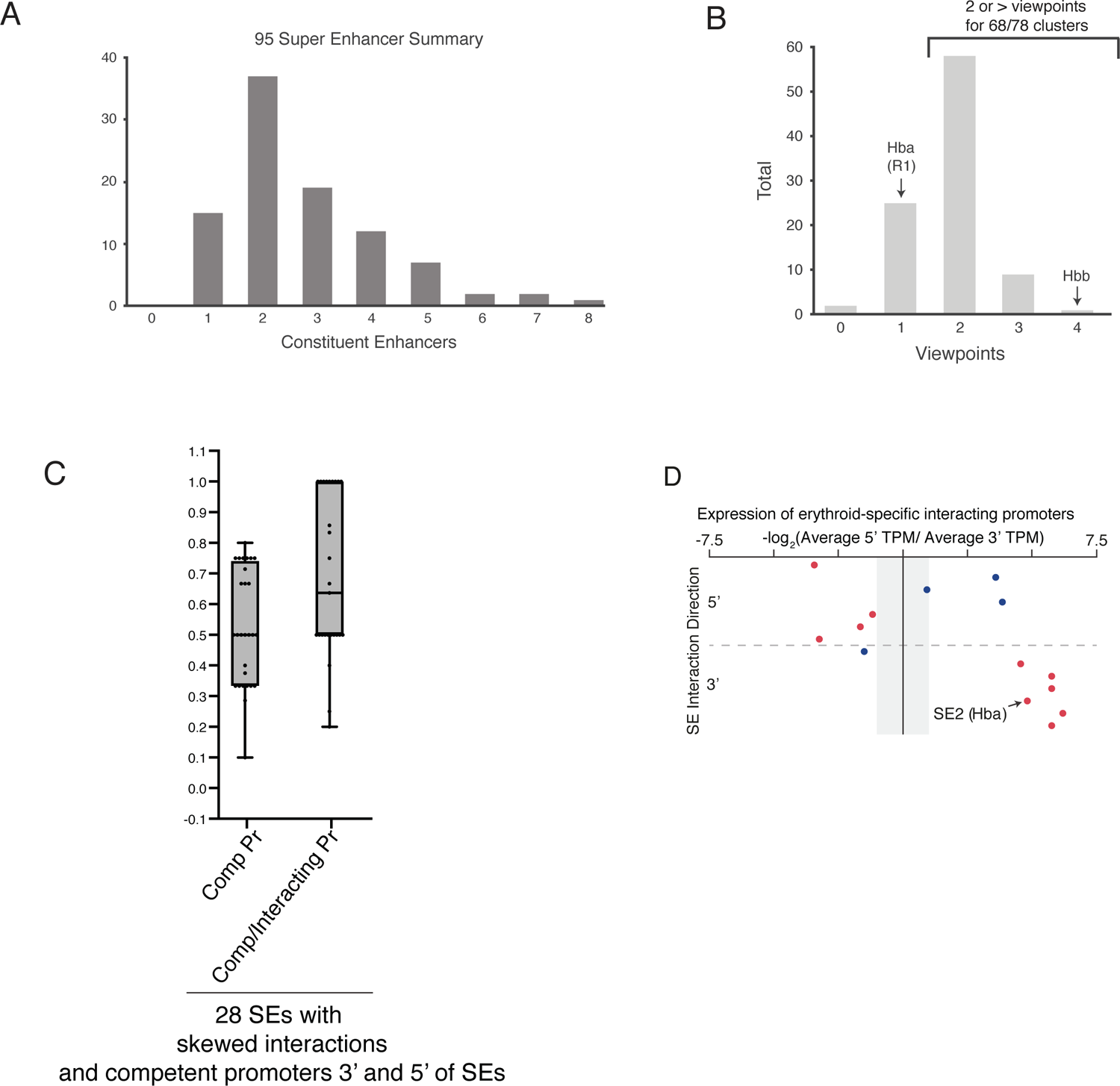
Description of the 95 erythroid SEs studied using NG Capture-C. (A) Based on the ROSE algorithm and enhancer element identification, the 95 erythroid SEs are composed of varying numbers of constituent enhancers as shown; 15 comprise 1 enhancer-like element and the rest contain two to 8 enhancer-like elements. (B) The distribution of NG Capture-C viewpoints across the SEs; designs for two SEs failed (indicated as zero on the x-axis), 1 viewpoint was designed for the single-enhancer SEs (15) and 10 of the multipartite SEs. The remaining 68 multipartite SEs were captured with at least two viewpoints. (C) The physical distribution of competent promoters relative to their associated SEs. As in Fig 1D, except this dataset represents SE in which competent promoters (H3K4me3+) are found on both sides of the relevant SEs (28 SEs). The first box plot (Comp Pr) accounts for all such competent promoters and shows an even distribution (mean 0.5). The second box plot (Comp/Interacting Pr) shows a skew (mean 0.7) when H3K4me3+ promoters are also selected for their differential interaction in erythroid cells. A ratio above 0.5 indicates that both the distribution of such promoters and the interaction skew are in the same direction indicating that the SEs are interacting with competent promoters in a directional manner. (D) Relating the direction of SE interactions to gene expression: as in Fig 1E, except this subset of interaction and expression analyses is done on SEs that are flanked on both sides by competent promoters with an erythroid-specific interaction profile (14 SEs). The horizontal dashed grey line represents the separation between the SEs whose interactions are skewed to upstream (5’) (above the line) or downstream (3’) (below the line) of the SE. The vertical axis separates relative fold difference in nascent expression (transcripts per million, TPM) of the erythroid-specific genes associated with their corresponding SEs. The grey shaded region on the y-axis indicates less than 2-fold change in expression. Red dots mark the SEs where the interaction direction skew and increase in levels of expression of the erythroid-specific genes are concordant. SE2 (*Hba* locus) is indicated (black arrows).

**Figure S2.**
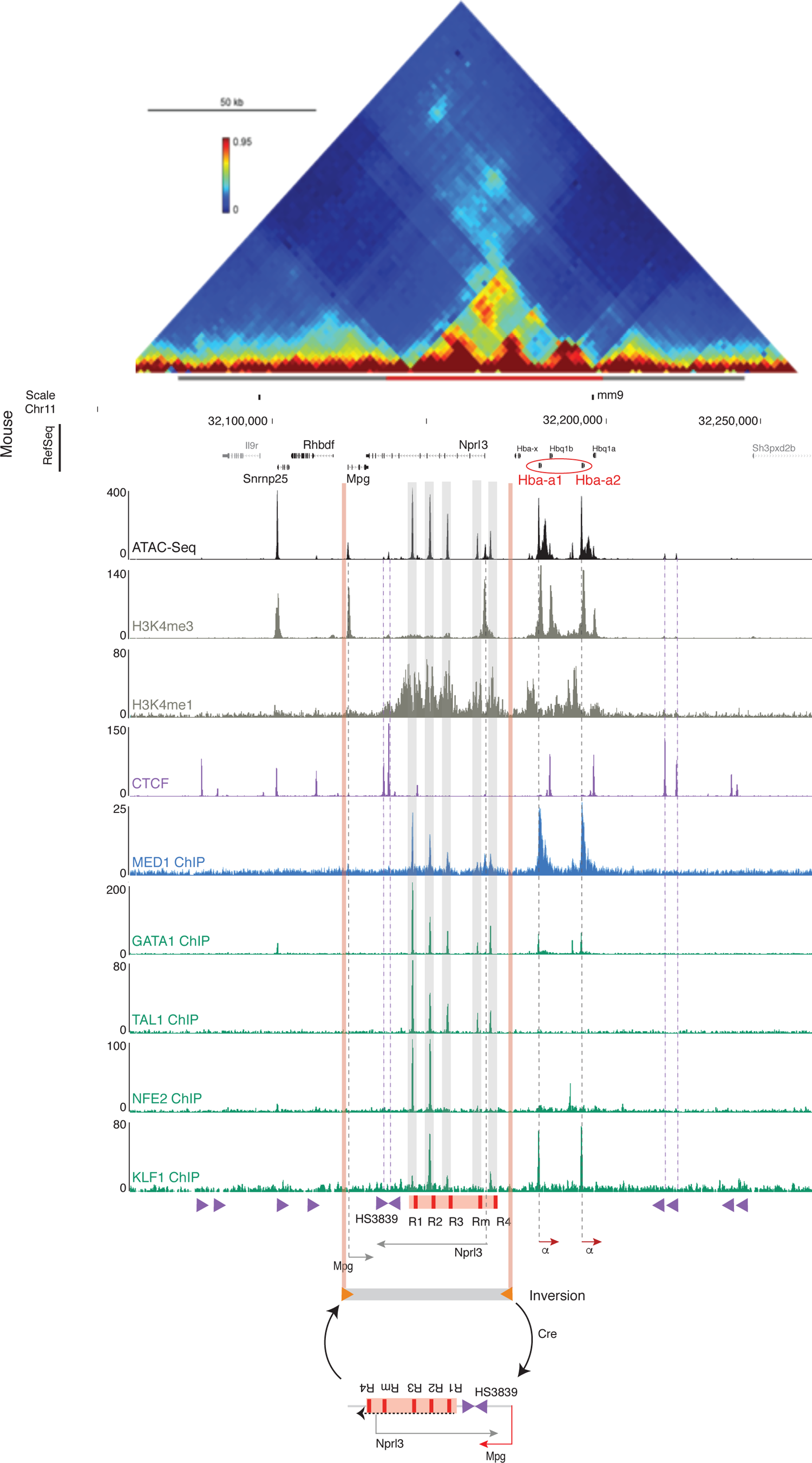
The well-characterised α-globin locus, a model to study mutipartite SE functional polarity. From top to bottom: 3C contact matrix of 200 kb spanning the mouse α-globin cluster; accessible chromatin (ATAC-seq); histone modifications (H3K4me3 and H3K4me1); occupancy of specialised and erythroid transcription factors (CTCF, MED1, GATA1, TAL1, NFE2 and KLF1). The grey and red bars immediately below the contact matrix represent the TAD (chr11:32,080,000-32,245,000) and sub-TAD (chr11:32,136,000-32,202,000), respectively. The α-globin genes are marked in red and the enhancers are highlighted in grey shaded vertical bars. Below the tracks, a schematic representation of the main regulatory elements spanning the α-globin locus, marked by dashed grey and purple vertical lines and indicating the adult α-globin genes (α) and flanking genes (*Nprl3, Mpg*) in red and grey arrows pointing in the direction of their expression and the CTCF-binding sites and their corresponding orientation in purple arrows, with the α-globin tested 5’ boundary elements labelled (HS3839). Orange box represents the SE and the vertical red bars mark the five constituent enhancers (R1-R4). Orange shaded vertical bars mark the limits of the inversion encompassing the SE as well as the *Nprl3* and *Mpg* genes, an interval that was flanked by convergent heterotypic LoxP sites (grey bar flanked by orange convergent arrows) and was flipped upon expression of Cre-recombinase in mESCs, as shown in the schematic.

**Figure S3.**
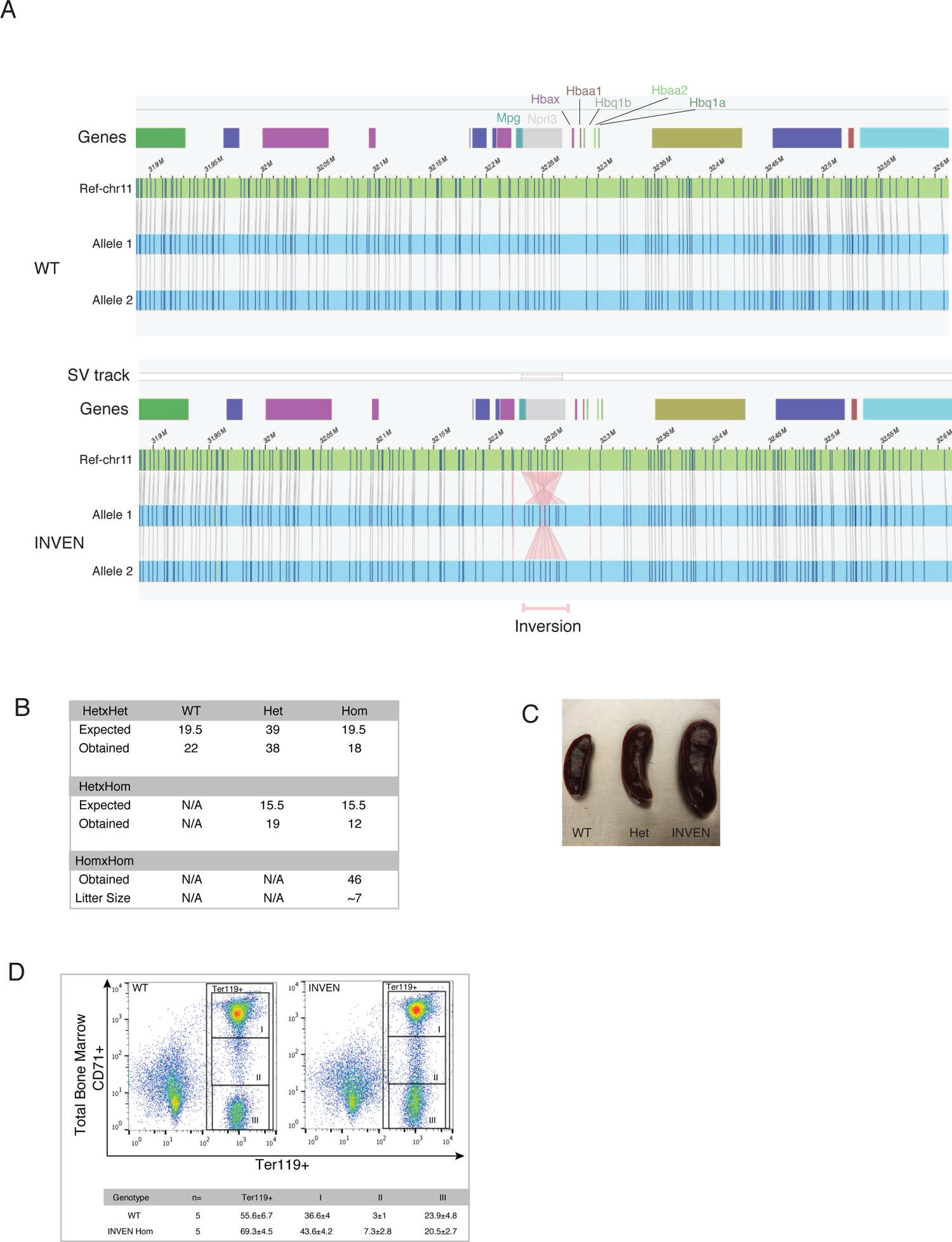
INVEN allele integrity confirmed by optical mapping and homozygous mice are viable with a phenotype reminiscent of stress erythropoiesis (A) Optical mapping using Bionano shown within ∼800Kb window (mm10 chr11:32,880,000-32,600,00) detects the inversion as a structural variant (SV track, marked in a pink box) and otherwise intact genomic region. No other genetic changes were picked up genome-wide. Genes are shown at the top of the tracks as coloured boxes with annotated genes within the α-globin locus. Ref-chr11 is the wildtype reference sequence provided by the Bionano Saphyr software. Allele 1 and allele 2 indicate both alleles in WT and INVEN samples. DNA extracted from wildtype and homozygous INVEN mouse ESCs (clone A4.2. This clone was derived from the heterozygous mESC clone injected in blastocysts to produce the mouse INVEN model. The INVEN homozygous A4.2 clone was also the base clone from which the other INVEN-based genetic models derived from. (B) The observed number of mice resulting from heterozygote (HetxHet) and heterozygote-homozygote (HetxHom) crosses fulfil the expected mendelian ratio (25:50:25). Live homozygote mice confirm no embryonic lethality in INVEN phenotype. (C) A gradual expansion of spleen is observed from WT to homozygous INVEN mice. (D) Immunophenotyping using red cell specific surface markers (Ter119 and CD71) show an expansion of the early erythroid compartments (I and II) in INVEN-derived bone marrow from adult mice that have not been exposed to any chemical treatment. No perturbation of the mature red cell compartment (III) in INVEN-derived bone marrow compared to that from WT.

**Figure S4.**
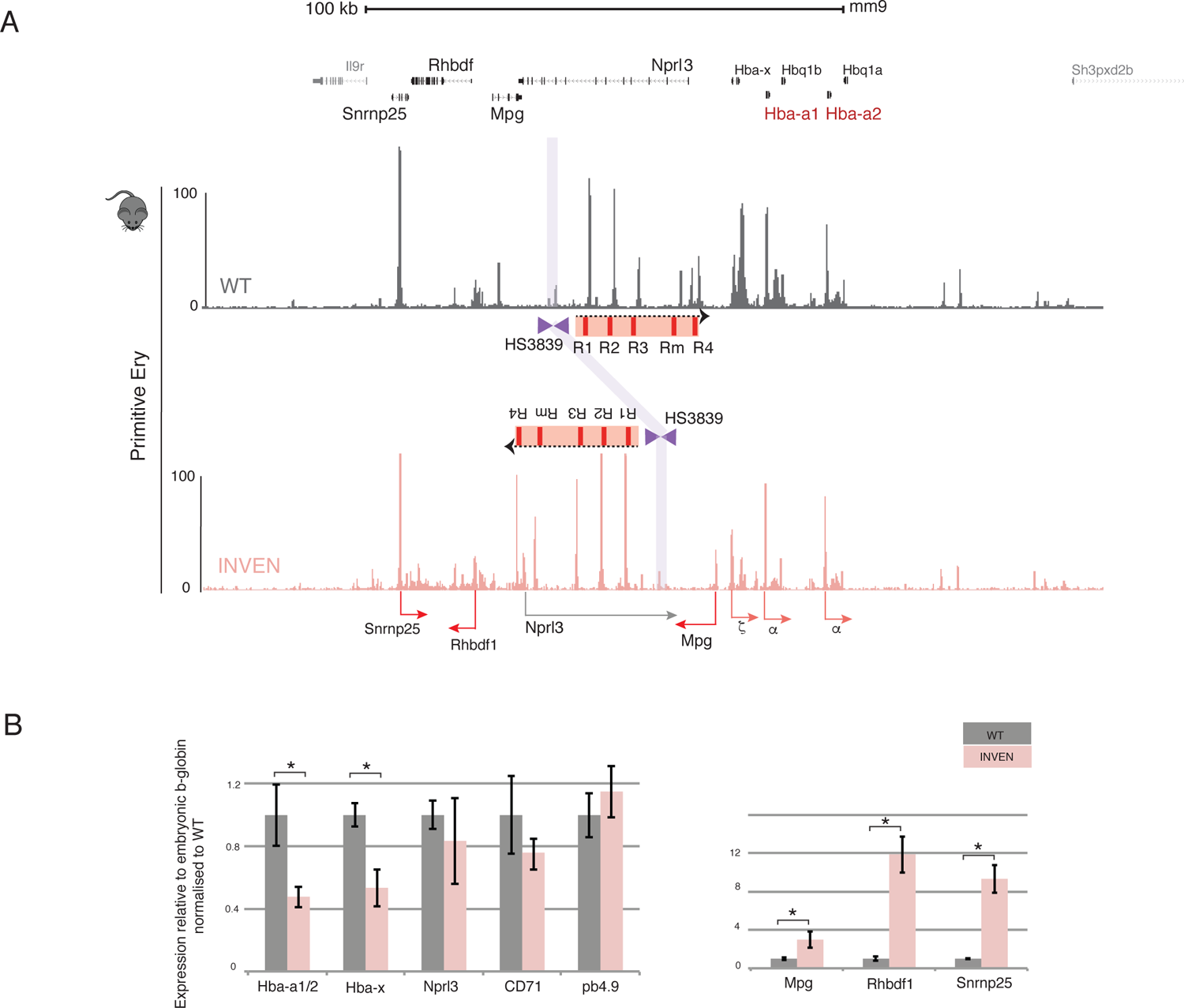
Gene Expression is perturbed at the α-globin locus in the INVEN E10.5 primitive erythroid cells. (A) ATAC-seq tracks show chromatin accessibility profiles in primary primitive erythroid cells derived from WT (grey) and INVEN (pink) E10.5 embryos. Note the differences in open chromatin corresponding to the genes in the locus. (B) Expression analysis by Real-time qPCR assessing level of mRNA expression for genes of interest relative to the embryonic beta globin and WT (as in Fig4C). Three embryos were analysed from each genotype. *P* values are obtained via an unpaired, two-tailed Student’s t-test. * *P* <0.0001.

**Figure S5.**
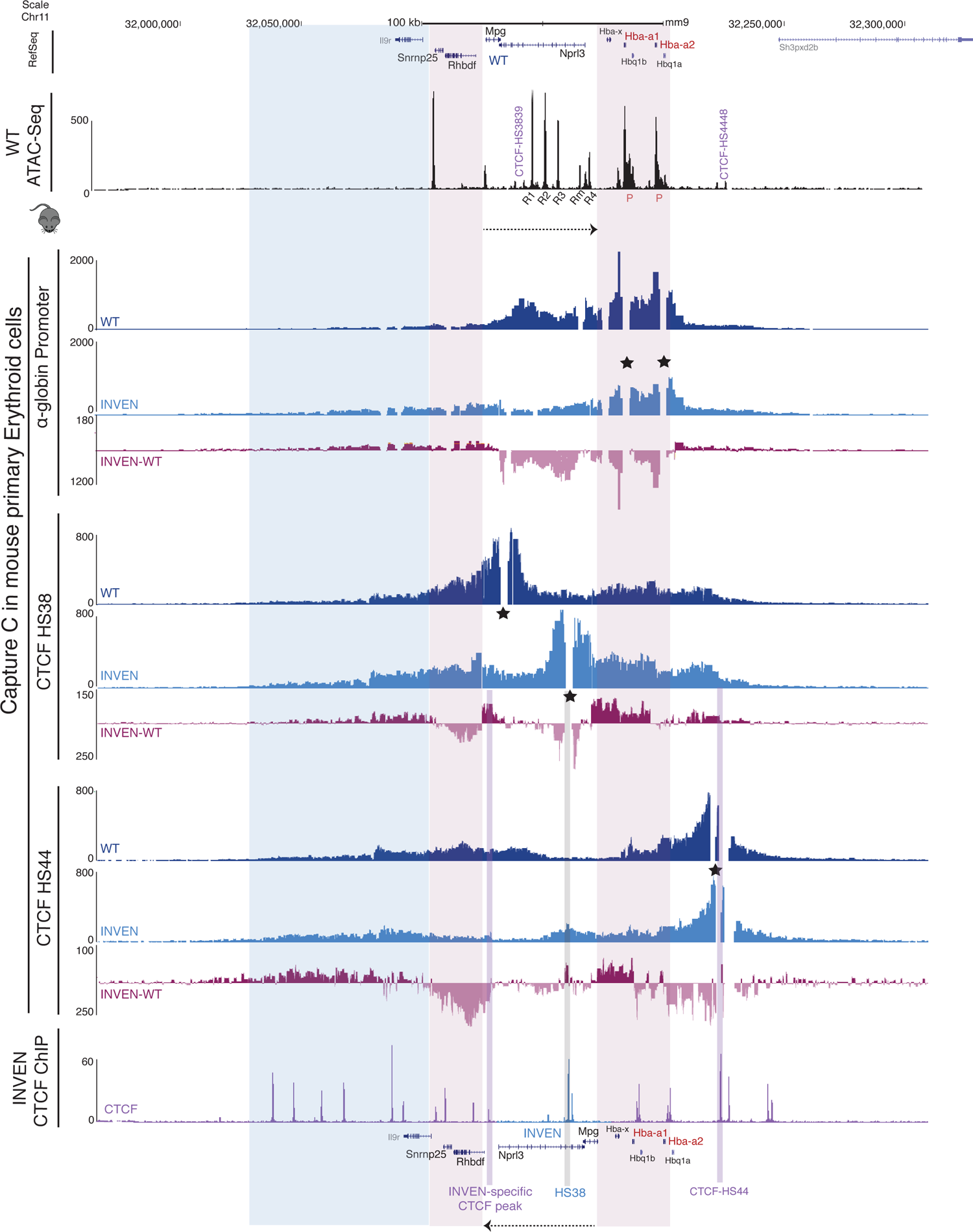
Comparison of NG-Capture C profiles shows redirected interactions from promoters and CTCF sites between WT and INVEN primary definitive erythroid cells across the α-globin locus. At the top, scale and RefSeq gene annotation and the bottom the same gene annotation but in the inverted configuration. All the regulatory elements (promoters P, enhancers R1-R4, and CTCF sites) are marked. Top and bottom tracks show normalised (reads per kilobase per million mapped reads, RPKM) and averaged read densities from 3 independent experiments of WT ATAC-seq and INVEN CTCF ChIP-seq, highlighting open chromatin and CTCF occupancy in primary definitive erythroid cells derived from WT and INVEN mice respectively. The NG Capture-C interaction profiles in WT (navy blue) and INVEN (light blue) show means of interacting fragment counts (n=3 independent biological replicates) using a 6 kb window. Additional track shows subtraction (INVEN-WT) per *Dpn*II fragment of significantly interacting fragments using DESeq2 (p.adj<0.05) with light pink for reduced interactions and dark pink for increased interactions in INVEN erythroid cells. The dashed black arrows indicate the direction of the SE in both WT and INVEN models. The stars mark the viewpoints used in the NG Capture-C experiment: the α-globin promoters, 5’ CTCF H38 site highlighted in a shaded blue bar in its new position in INVEN, and 3’ CTCF HS44 site and the INVEN-specific CTCF peak highlighted in shaded purple bar. The shaded pink and blue boxes indicate regions of changed interaction profiles in the INVEN model compared to WT.

**Figure S6.**
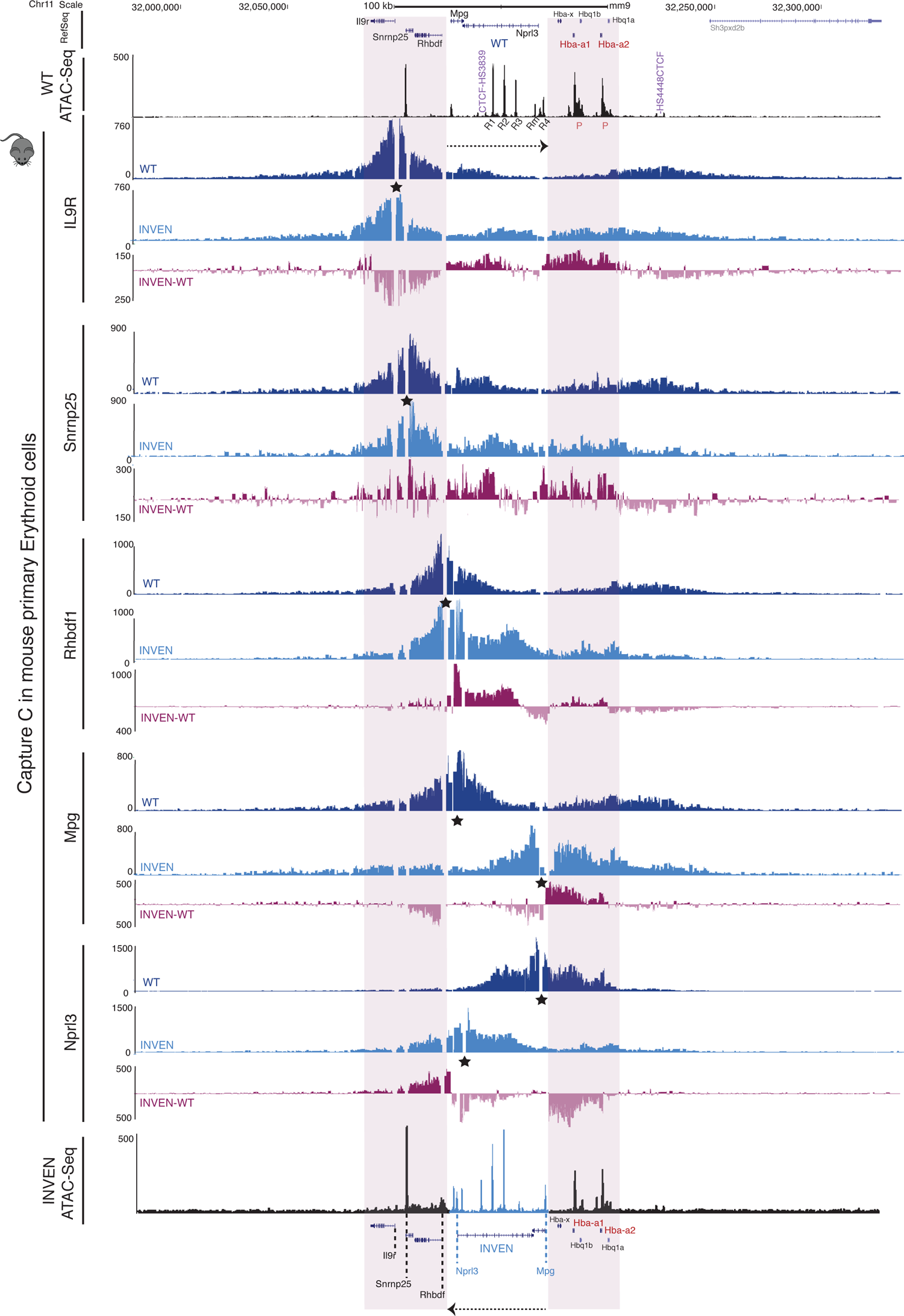
NG-Capture C profiles show redirected interactions from all the promoters present within across the α-globin locus when comparing WT and INVEN primary definitive erythroid cells. Same as in Fig 6 when the annotations are marked in the figure, except that the bottom track is of an ATAC-seq for the INVEN model and the NG Capture-C viewpoints, as indicated by the stars above the corresponding peaks highlight the promoters of the genes *IL9R, Snrnp25, Rhbdf1, Mpg, Nprl3*, from top to bottom. Note the gain and loss of interactions across the α-globin locus.

**Figure S7.**
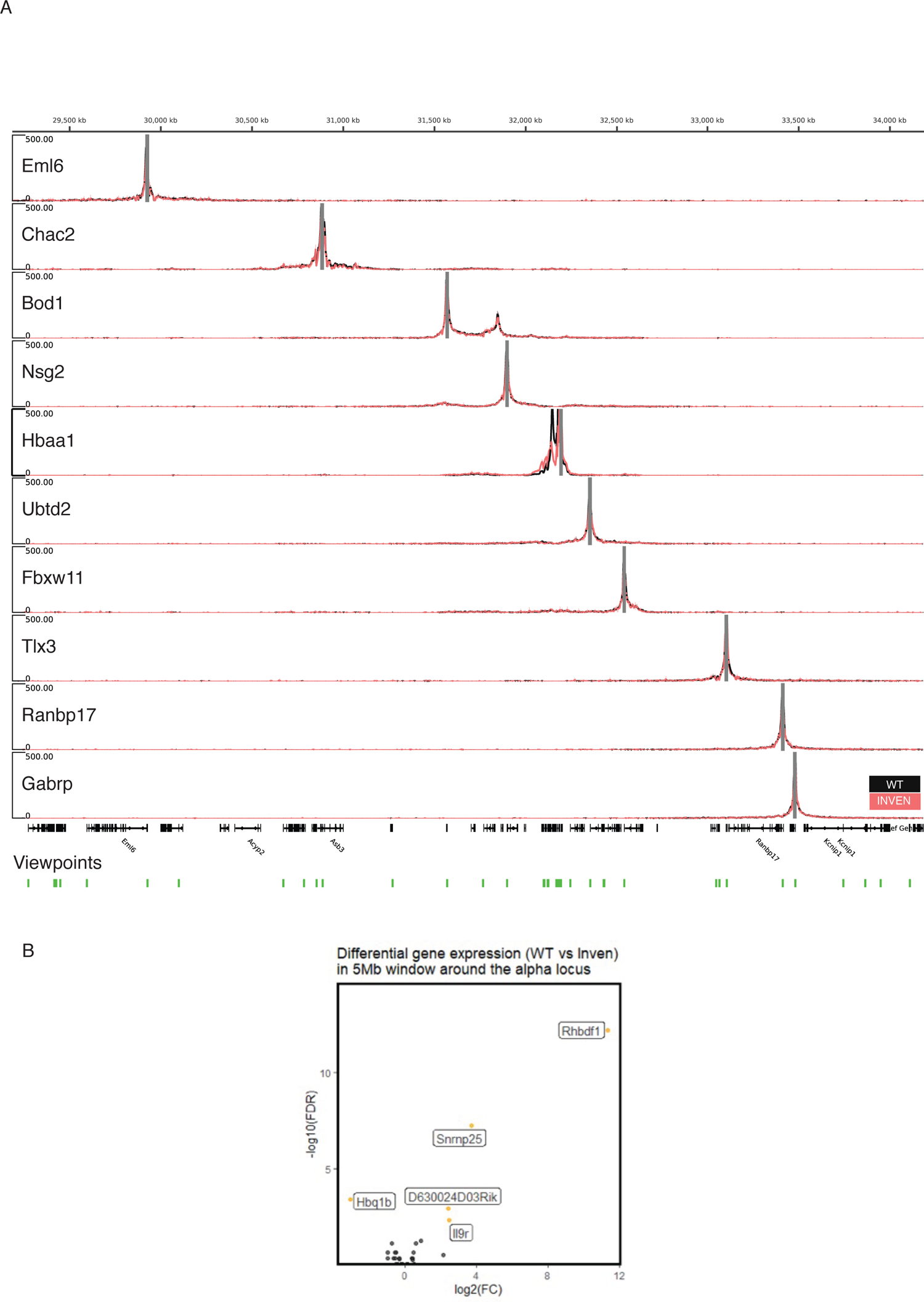
NG-Capture C profiles show undisturbed interactions from all the promoters captured within 5.6Mb flanking the α-globin locus when comparing WT and INVEN primary erythroid cells. (A) Windowed mean 3C interactions over 4.5 Mb (mm9 chr11:29500000-34000000) for 10 representative promoter NG capture-C viewpoints as indicated captured with a pool of 42 viewpoints for all the promoters spanning 5.6Mb flanking the α-globin locus. Domain interactions are unaffected in INVEN compared to WT except at the Hbaa2 viewpoint (other affected promoters also shown in Fig S6). Solid lines show means (n=3 independent 3C libraries for each of WT and INVEN primary erythroid cells derived from spleen of adult mice) with one standard deviation (shading) using a 6 kb window. (B) Differential expression between WT and INVEN primary erythroid cells within the 5.6Mb region flanking the alpha globin locus; orange dots represent genes with > than 2-fold expression change. The only genes affected within the 5.6Mb are the ones contained in the across the α-globin locus. PolyA+ RNA-seq data extracted from the main data set presented in Fig 5D.

**Figure S8.** Deletion of the 5’ α-globin boundary element (HS3839 CTCF) in INVEN mESC model does not rescue the INVEN phenotype in *in vitro* derived erythroid cells. At the top, scale and RefSeq gene annotation. Normalised (reads per kilobase per million mapped reads, RPKM) and averaged read-densities from 3 independent experiments of ATAC-seq show open chromatin in EB-derived erythroid cells differentiated from WT and INVEN mESCs. NG Capture-C interaction profiles in WT (grey), INVEN (pink), CTCF-KO (light blue) and INVEN-KO (purple) show means (n=3 independent biological replicates) of interacting fragment count using a 6 kb window. Additional track shows subtractions (INVEN-WT, (INVEN)-(CTCF-KO), (INVEN-KO)-(INVEN)) per *Dpn*II fragment of significantly interacting fragments using DESeq2 (p.adj<0.05) with light pink for reduced interactions and dark pink for increased interactions across the models analysed. The dashed black arrows indicate the direction of the SE in both WT and INVEN models. The star marks the viewpoint (the R1 enhancer) used in the NG Capture-C experiment. The purple shaded bar indicates *Mpg* gene, located between the SE and the α-globin genes in the INVEN models. Note the increased interactions at the newly positioned *Mpg* in the INVEN-KO. The dashed black arrows indicate the direction of the SE in both WT (top) and INVEN (bottom) models. The stars mark the viewpoint (R1 enhancer) used in the NG Capture-C experiment. The purple shaded box indicates the gained interaction over the *Mpg* gene in INVEN-KO erythroid cells.

**Figure S9.** NG-Capture C profiles show redirected interactions from all the promoters present within the α-globin sub-TAD when comparing α-globin 5’ boundary (HS3839) knock out (CTCF-KO) and HS3839 knock out in INVEN model (INVEN-KO) in EB-derived erythroid cells. Same as in Fig S6 except; the data is from CTCF-KO (HS3839 deleted in WT) and INVEN-KO (HS3839 deleted in INVEN) EB-derived erythroid cells.

## Methods

### Mice

All mouse work was performed in accordance with UK Home Office regulations under project license number 30/3339 with appropriate local ethical review. All animals were housed in Individually Ventilated Cages with enrichment, provided with food and water ad libitum, and maintained on a 12 h light: 12 h dark cycle (150-200 lux cool white LED light, measured at the cage floor). Mice were given neutral identifiers and the haematological phenotype was analysed by research technicians unaware of mouse genotype during outcome assessment. Samples for ATAC-seq, ChIP-seq, NG Capture-C, and gene expression analyses were not randomized and the investigators were not blinded to allocation during these experiments and outcome assessments.

The INVEN mouse model was generated by blastocyst injection of targeted E14Tg2a mouse Embryonic Stem Cells (mESCs) heterozygous for the inversion of the α-globin super-enhancer (SE) into a C57BL/6J background mice. The inversion is described in Fig 2B and S2.

### Isolation of erythroid cells derived from adult mouse spleen

Primary Ter119+ erythroid cells were obtained from the spleens of adult mice that were treated with phenylhydrazine as described previously^1^. Spleens were mechanically dissociated into single cell suspensions in cold phosphate-buffered saline (PBS; Gibco: 10010023)/10% fetal bovine serum (FBS; Gibco: 10270106) and passed through a 70 μm filter to remove clumps. Cells were washed with cold PBS/10% FBS and resuspended in 10 μl of cold PBS/10% FBS per 10^6^ cells and stained with a 1/100 dilution of anti-Ter119-PE antibody (BD Pharmingen 553673 2 μg/ml) at 4 °C for 20 minutes. Stained cells were washed with cold PBS/10% FBS and resuspended in 8 μl of cold PBS/0.5% BSA/2 mM EDTA and 2 μl of anti-PE MACS microbeads (Miltenyi Biotec: 130-048-801) per 10^6^ cells and incubated at 4 °C for 15 minutes. Ter119+ cells were positively selected via MACS lineage selection (LS) columns (Miltenyi Biotec: 130-042-401) and processed for downstream applications. Purity of the isolated erythroid cells was routinely verified by Fluorescence-Activated Cell Sorting (FACS).

### Hematological analysis of mouse models

Hematologic parameters for adult mice from INVEN model: WT, heterozygote and homozygote littermates were assessed where possible. A balanced mix of male and female were examined at more than 7 weeks of age to minimise between-individual variation. All mice were generated on the same complex background. Blood was collected into heparinized capillary tubes and blood counts were assayed using the Horiba Medical SciI Vet abc Plus+ instrument. Sample sizes were chosen with a view to detecting 25% reduction in hemoglobin concentration. Blood smears were prepared using 2 μl of peripheral blood spread on a slide, dried, stained, and mounted. The blood smears were either used for the reticulocyte counts performed manually after Brilliant Cresyl Blue (BCB) staining or for general red blood cell morphology assessment after May–Grünwald–Giemsa staining. Reticulocyte counts and blood smear assessment were performed by two independent assessors blinded to genotype.

### Genome engineering to generate the genetic models analysed in this study

#### Mouse Embryonic Stem Cells (mESCs) targeting strategy to generate the inversion of the *α-globin* SE (INVEN model)

The region to invert encompasses the SE but was delimited such that the breakpoints do not impinge on the integrity of the locus such as the transcription of *Nprl3* and *Mpg* housekeeping genes. The strategy also ensured that the chromatin at the boundaries of the inversion are not marked by any histone modifications or bound by any of the factors assessed (Fig S2), to avoid disruption of a potentially functional DNA region.

The strategy was based on a similar approach (Recombination-Mediated Genome Replacement or RMGR) whereby the mouse α-globin locus was replaced by the human α-globin locus^2^. In outline, two targeting vectors, each harbouring a LoxP site flanked by sequences homologous to either the 5’ or the 3’ insertion sites at the α-globin locus, were assembled following a recombineering strategy in E.Coli (Methods Fig 1A). The vectors were then electroporated into mESCs for a sequential insertion of the 3’ and then 5’ convergent LoxP sites at the α-globin locus (insertion sites at the edges of this interval, chr11:32,224,538-32,271,838). Positive resistance markers in the 3’ and 5’ targeting constructs (Hygromycin and Neomycin respectively) secured the selection for LoxP integration along with an incomplete 5’ or 3’ components of an Hprt minigene at the flanks of the ∼48Kb sequence. Sequential employment of site-specific recombinases induces the inversion of the 48Kb sequence (Cre), and deletion of all selectable markers (Flp) (Methods Fig 1B-C). Upon inversion induced by Cre, selection is done by HAT-resistance only in mESCs where the integration of both targeting vectors occurred *in cis* and resulted in the complementation of the Hprt gene (3’ and 5’ parts of the gene). Any other events would be result in nonviable mESCs (dicentric chromosomes).

Southern blots and PCR strategies were used for the screening for targeted events both at the 3’ and 5’ integration sites (data available upon request). Table 1 contains primers for the Southern blot probe amplicons.

The strategy and data for retargeting clones heterozygous to INVEN (WT/INVEN) following Hprt minigene excision and then excision of the Hprt minigene in the resulting homozygous INVEN/INVEN clones (A4.2, B4.4, F11.2) is shown in (Methods Fig 2A, B, C, D).

The integrity of the INVEN homozygous clone (A4.2) was checked using an orthogonal method; optical mapping using Bionano Saphyr technology.

### Optical mapping using Bionano Saphyr technology

High molecular weight DNA from patient derived lymphoblastoid cell lines was isolated using the Bionano plug lysis protocol, using the Bionano Prep Blood and Cell Culture DNA isolation kit. Genomic DNA was barcoded using the Bionano Prep DLS labelling kit, in accordance with the manufacturer’s instructions. Labelled samples were analysed using the Saphyr device, and optical maps were assembled and structural variants analysis was conducted using Bionano Access software.

### CRISPR-Cas9 strategies and mESC transfections

We used CRISPR-Cas9 targeting strategies for the generation of various mESC models used in this study.

For generation of enhancer deletion R1 (DelR1) and R2 inversion (INVR2) mESCs, a similar strategy was followed; Two sgRNAs that flank the enhancer sequence were designed and cloned into a pSpCas9(BB)-2A-GFP (pX458) vector, a gift from Feng Zhang (Addgene plasmid #48138; http://n2t.net/addgene:48138; RRID:Addgene_48138) or a modified vector with the GFP tag exchanged for an mRuby cassette (pX458-Ruby). For generation of HS3839 deletion (D3839) mESCs, sgRNAs were designed to flank a sequence encompassing the two CTCF sites (at HS-38: TACCCTCTGGTGGC and at HS-39.5: CTGGCCACTGGGGG separated by around 1.5kb). Four guides for a CRISPR/Cas9 nickase strategy were cloned into p335 (Cas9-D10A) modified vector containing a neomycin selectable marker. An HDR vector of 2991 bp was designed such that a fragment encompassing both CTCF sites and the intervening 1447bp sequence was flanked by homology arms of 750bp. sgRNA recognition sequences outside the homology arms were also included and resulted in the release of the HDR sequence from the donor vector upon co-transfection with sgRNA/Cas9 plasmids. Protospacer Adjacent Motif (PAM) sequences were mutated where necessary to avoid cutting the HDR plasmid. CTCF consensus sequences were replaced by ScaI sites (AGTACT) for ease of screening. Modified sequences were checked using Sasquatch tools for undesirable creation of potentially active hypersensitive sites in erythroid cells^3^. Similar strategy was followed to produce MpgKO except that HDR was achieved using a ssODN of 200bp with 97 and 94bp homology with the flanks of *Mpg* exon1 and a ScaI restriction site (bold) inserted in the middle for screening purposes.

ssODN: TCTGTGGCCCACCCTATCCCAAGGCAGAGCCTTCTGGTCCGGGACCCTTGAAAATTCA GTCCGGGTTCAGAGAGGATGTCTTGATGGCAATCGGGCC**AGTACTT**GGACGCAGGAA AATCAGGAGTTCAAAGCCAGCATAGGCCACATGGGACCCTGTCCTTTAAAAGAGTAATGGCTGTTTAACTAGTCAGCTCAGC

To generate DelR1, INVR2, and CTCFKO, mESC transfections were performed using the Neon electroporation system (Invitrogen), according to the manufacturer’s instructions. A transfection protocol optimized for mESCs (3 pulses of 1400V/10ms) whereby 10^6^ mESCs were resuspended in buffer R and electroporated with a total of 5 μg plasmid DNA (2.5ug/gRNA for DelR1 mESCs,) and 11ug (1.25ug/gRNA and 6ug for HDR vectors for INVR2 and D3839). After electroporation, DelR1 and INVR2 targeted cells were cultured for 48-72 hours prior to FACS sorting the eGFP-Ruby expressing population, which was then seeded at clonal density on gelatinized 10cm dishes for at least 6 days. D3839 cells were directly plated on a gelatinized 10cm dish at clonal density and targeted clones were selected for neomycin resistance (G418 at 200μg/ml) 24h to 72h post transfection and left to grow for at least 6 days. Colonies were picked into 96-well plates and screened for mutations by PCR and Sanger sequencing. For MpgKO generation, mESCs were transfected using the Fugene 6 Transfection Reagent (Promega), according to the manufacturer’s instructions. In brief, 4ug of total DNA (1ug for each of sgRNA and 2ug of ssODN HDR donor) were incubated with 20ul Fugene 6 reagent/180 ul media for about 15min before being added to 10^6^ cells of mESCs in a well of a 6-well plate. After 48hrs, the mESCs were FACS sorted for GFP and Ruby, the reporters of the sgRNA vectors. Sorted cells were plated at clonal density (2000 cells/10cm dish) and colonies picked (2×96 well plates/mESC clone transfected) and screened using a restriction digest insertion; PCR the fragment across the expected deletion/insertion followed by ScaI digestion.

For sgRNA and screening PCR primer sequences, see Tables 2, 3.

### mESC in vitro hematopoietic differentiation; Embryoid Body (EB) differentiation

24-48 h prior to differentiation, cells were induced by passaging into base media (Iscove’s modified Dulbecco’s medium (IMDM), 1.4×10^-4^ M monothioglycerol (Sigma-Aldrich) and 50 U/ml penicillin-streptomycin (Thermo Fisher)) supplemented with 15% heat-inactivated fetal calf serum (ΔFCS, Thermo Fisher) and 1000 U/ml LIF.

For EB generation, cells were disaggregated by trypsinisation and quenched in base media (as above) supplemented with 10% ΔFCS. Differentiation media was prepared fresh on the day of differentiation by supplementing base media (as above) with 15% ΔFCS, 5% protein-free hybridoma medium (PFHM-II, Thermo Fisher), 2 mM L-glutamine (Thermo Fisher), 50 μg/ml L-ascorbic acid (Sigma Aldrich), 3×10^-4^ M monothioglycerol and 300 μg/ml human transferrin (Sigma Aldrich). Cells were plated in petri dishes (Thermo Fisher) at 1.5-3×10^4^ cells in 10 ml differentiation media. EBs were left to differentiate for up to seven days without disruption except for gentle manual shaking every few days to disrupt potential EB aggregation and/or attachment to the dishes.

EBs from 10 cm dishes were harvested by collection of the entire plate contents into falcon tubes, spinning at 1000 rpm for 5 minutes before and after a single PBS wash step to ensure complete recovery. After PBS removal, EBs were disaggregated in 0.25% trypsin (0.3 ml per 10 cm dish) for 3 minutes at 37°C with continuous manual shaking to prevent sedimentation at the bottom of the tube. Trypsin was quenched with an equal volume of FCS and a single-cell suspension obtained through tituration. Cells were collected by spinning at 1200 rpm for 5 minutes and resuspended as needed.

### Erythroid population isolation from EBs

CD71+(high) cells were isolated by magnetic column separation (LS Column, Miltenyi), according to the manufacturer’s instructions as previously described^4^. Briefly, cells were labelled with anti-mouse CD71-FITC (eBioscience 11-0711-85; 1:200) in staining buffer (PBS with 10% FCS; 500 μl per 10^7^ cells) for 20 minutes at 4°C, rolling, then washed by adding staining buffer (1 ml per 10^7^ cells) and spinning. After supernatant removal, cells were incubated with MACS anti-FITC separation microbeads (Miltenyli; 10 μl per 10^7^ cells) in ice-cold separation buffer (PBS plus 0.5% bovine serum albumin (BSA) and 2 mM EDTA; 90 μl per 10^7^ cells) for 15 minutes at 4°C, rolling, and washed by adding separation buffer (1 ml per 10^7^ cells) and spinning. Bead-labelled cells were resuspended in 500 μl cold separation buffer and added to a pre-equilibrated LS column. The negative fraction was washed through with two flushes of 3 ml cold separation buffer and the positive fraction collected by forcing cells from the column in 5 ml separation buffer. After spinning and supernatant removal, cells were resuspended in staining buffer as needed for downstream processing. Population purity and selection efficiency were determined by flow cytometry. Information on the antibodies used are in Table 4.

### ATAC-seq and ChIP-seq

Assay for Transposase-Accessible Chromatin (ATAC)-seq was performed on 75,000 Ter119+ cells isolated from phenylhydrazine-treated mouse spleens as previously described^5,6^. ATAC-seq libraries were sequenced on the Illumina Nextseq platform using a 75-cycle paired-end kit (NextSeq 500/550 High Output Kit v2.5: 20024906).

CTCF Chromatin immunoprecipitation (ChIP) was performed on 1 x 10^7^ Ter119+ erythroid cells using a ChIP Assay Kit (Millipore: 17-295) according to the manufacturer’s instructions. Cells were crosslinked by a single 10 min 1% formaldehyde fixation. Chromatin fragmentation was performed with the Bioruptor Pico sonicator (Diagenode) for a total sonication time of 4 min (8 cycles) at 4°C to obtain an average fragment size between 200 and 400 bp. Immunoprecipitation was performed overnight at 4 °C with various antibodies (included in Table 5). Library preparation of immunoprecipitated DNA fragments was performed using NEBNext Ultra II DNA Library Prep Kit for Illumina (New England Biolabs: E7645) according to the manufacturer’s instructions. Libraries were sequenced on the Illumina Nextseq platform using either a 75-cycle paired-end kit (NextSeq 500/550 High Output Kit v2.5: 20024906) or a 300-cycle paired-end kit (NextSeq 500/550 Mid Output Kit v2.5: 20024905).

Data were analysed using an in-house pipeline^7^. PCR duplicates were removed, and biological replicates were normalised to Reads Per Kilobase per Million (RPKM) mapped reads using deepTools package^8^, averaged across three biological replicates using bedtools^9^, and visualised using University of California Santa Cruz (UCSC) genome browser^10^.

### NG Capture-C

Next-generation Capture-C was performed as previously described^11^. A total of 1-2 x 10^7^ Ter119+ erythroid cells were used per biological replicate. We prepared 3C libraries using the DpnII-restriction enzyme for digestion. We added Illumina TruSeq adaptors using the NEBNext Ultra II DNA Library Prep Kit for Illumina (New England Biolabs: E7645) according to the manufacturer’s instructions, and performed capture enrichment using NimbleGen SeqCap EZ Hybridization and Wash Kit (Roche: 05634261001), NimbleGen SeqCap EZ Accessory Kit v2 (Roche: 07145594001), and previously published custom biotinylated DNA oligonucleotides (R1 and HS-38 viewpoints^12^; α-globin promoters’ viewpoints^11^. NG Capture-C data were analysed using the CaptureCompendium toolkit^13^ which uses Bowtie^14^ to map reads to the mm9 mouse genome build. *Cis* reporter counts for each sample were normalised to 100,000 reporters for calculation of the mean and standard deviation (three biological replicates). Mean reporter counts were divided into 150 bp bins and smoothed using a 5 kb window.

### RNA expression analysis

Total RNA was isolated from 105-2×106 Ter119+ erythroid cells lysed in TRI reagent (Sigma-Aldrich: T9424) using a Direct-zol RNA MiniPrep kit (Zymo Research: R2050). DNase I treatment was performed on the column as recommended in the manufacturer’s instructions but with an increased incubation of 30 min at room temperature (rather than 15 min). To assess relative changes in gene expression by qPCR, cDNA was synthesised from 500ng-1 μg of total RNA using SuperScript III First-Strand Synthesis SuperMix for qRT-PCR (Invitrogen, ThermoFisher: 11752-050) according to the manufacturer’s instructions. The ΔΔCt method was used for relative quantification of RNA abundance using Fast SYBR Green Master Mix (Thermo Fisher) using the primers summarised in Table 2.10. Data were normalised to the 18S ribosomal gene or relevant β-globin genes using the ΔCT method, taking the mean of multiple control CT values when applicable. Primers used for the genes assessed are in Table 6.

For RNA-seq libraries, 1-2 μg of total RNA was depleted of rRNA and globin mRNA using the Globin-Zero Gold rRNA Removal Kit (Illumina: GZG1224) according to the manufacturer’s instructions. To enrich for mRNA, poly(A)+ RNA was isolated, strand-specific cDNA was synthesised, and the resulting libraries prepared for Illumina sequencing using the NEBNext Poly(A) mRNA Magnetic Isolation Module (New England Biolabs: E7490) and the NEBNext Ultra II Directional RNA Library Prep Kit for Illumina (New England Biolabs: E7760) following the manufacturer’s instructions. Poly(A)+ and Poly(A)-RNA-seq libraries were sequenced on the Illumina Nextseq platform using a 75-cycle paired-end kit (NextSeq 500/550 High Output Kit v2.5: 20024906). Reads were aligned to the mm9 mouse genome build using STAR^15^. DeepTools bamCoverage was used to calculate normalized (RPKM) and strand-specific read coverage, which was visualized in the UCSC genome browser. Mapped RNA-seq reads were assigned to genes using Subread featureCounts using RefSeq gene annotation. Normalized differential gene expression, between biological triplicate data from littermate wild-type and INVEN mutant mice extracted in parallel, was calculated with the DESeq2 R package.

## Data and code availability

All data generated for this study are included in this published article and its supplementary information. ChIP-seq, ATAC-seq, RNA-seq and NG Capture-C data (sequence reads and processed) is available in the Gene Expression Omnibus (GEO) under accession numbers XX. Codes used in the analysis of this manuscript are referenced.

## Methods tables

**Table 1.**
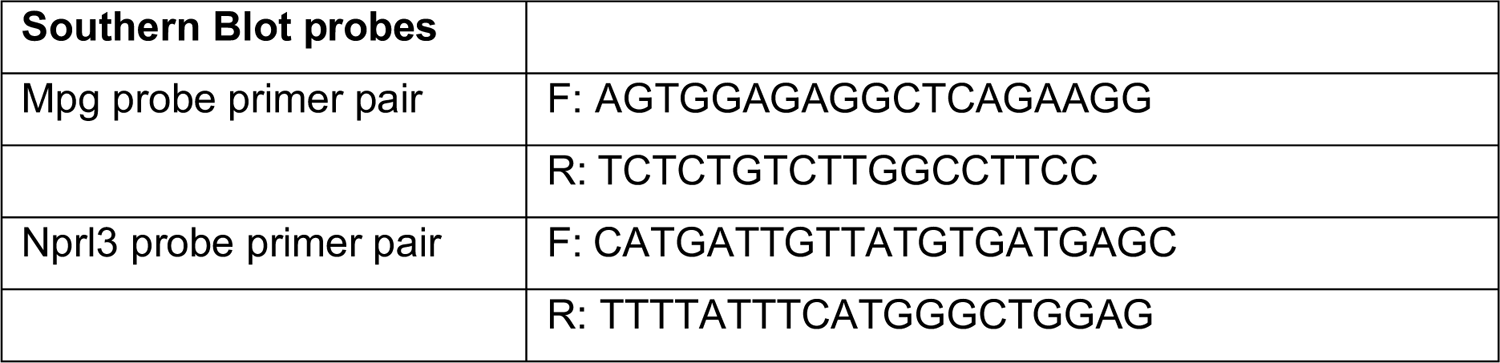
Sequences for Southern Blot probes and Sanger Sequencing primers used for screening INVEN models.

**Table 2.**
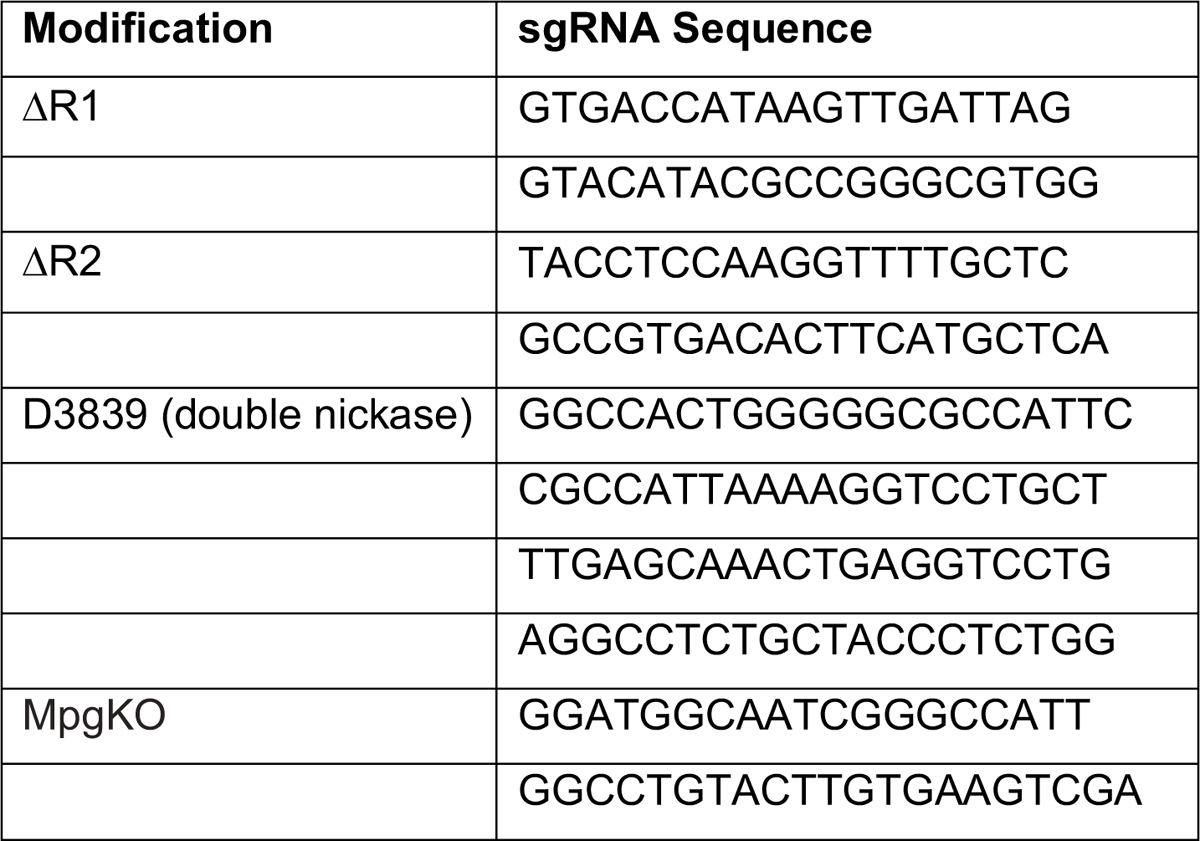
gRNA used for CRISPR-Cas9 genome editing.

**Table 3.**
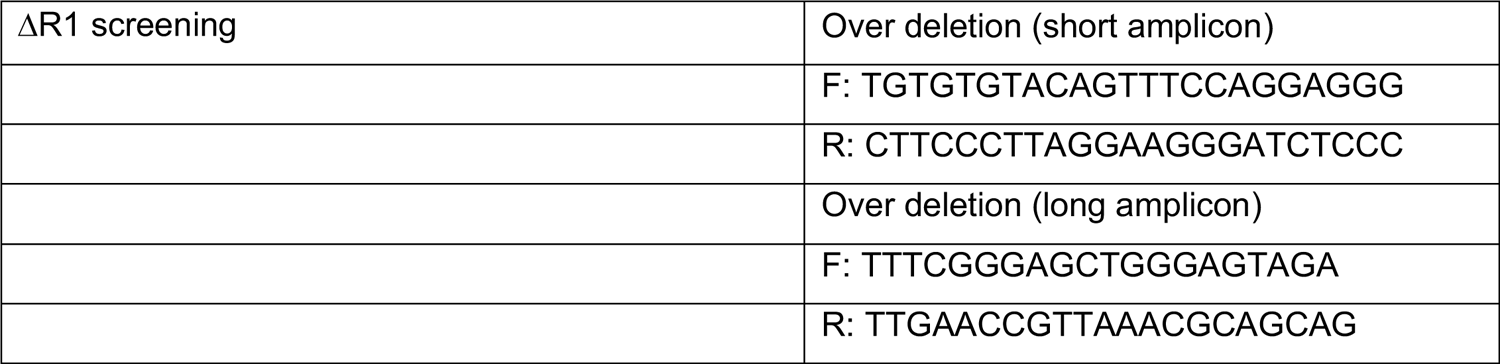
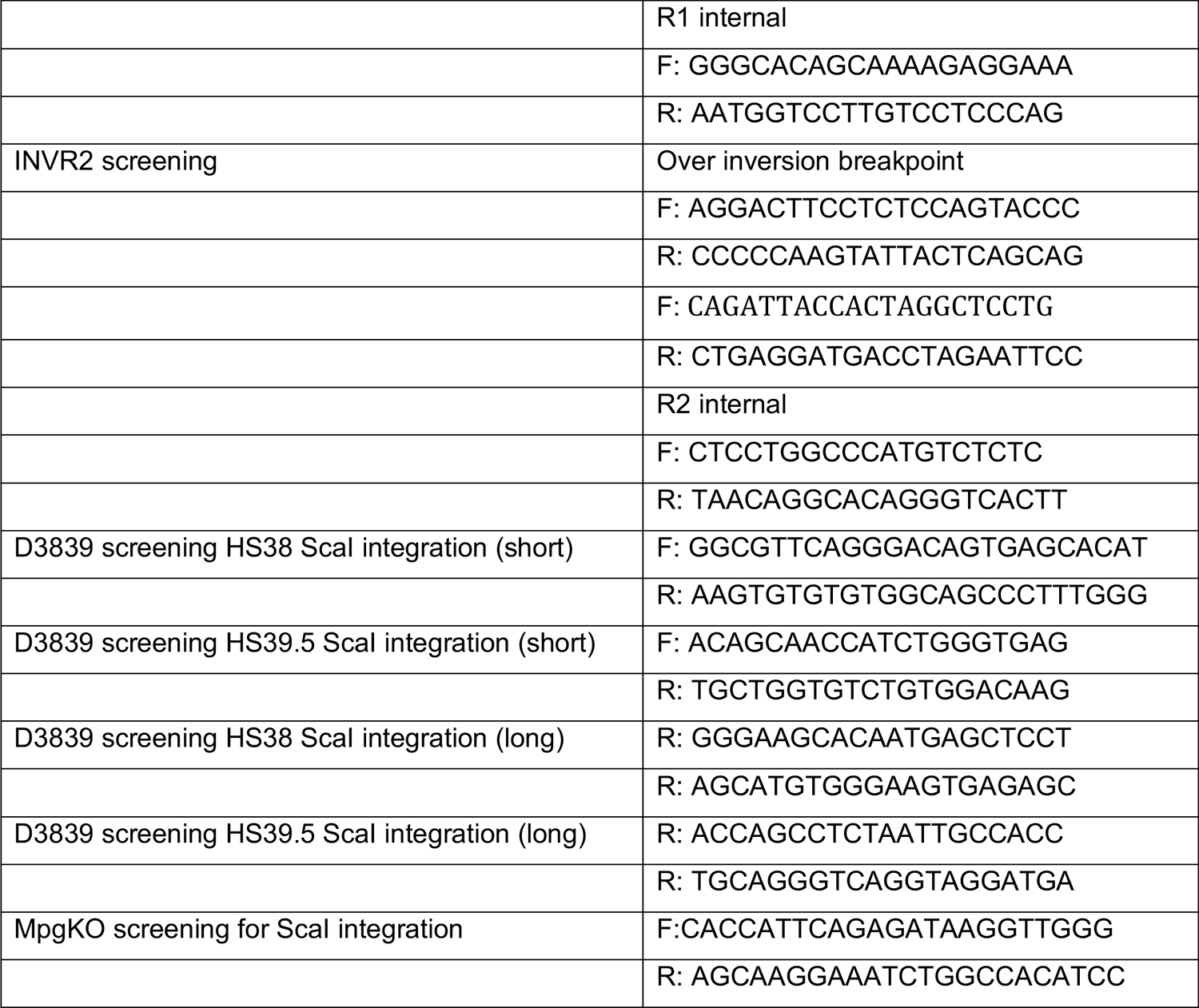
Primer pairs used for PCR screening of mESC targeted clones

**Table 4.** Flow cytometry and magnetic column purification antibodies.

| Protein target Antibody, source | Source | Working concentration |
| --- | --- | --- |
| FITC Rat Anti-Mouse CD71 | eBioscience 11-0711-85 | 2.5 µg/ml |
| PE Rat Anti-Mouse Ter119 | BD Pharmingen 553673 | 2 µg/ml |
| Hoechst | Invitrogen H3569 | 1 µg/ml |

**Table 5.**
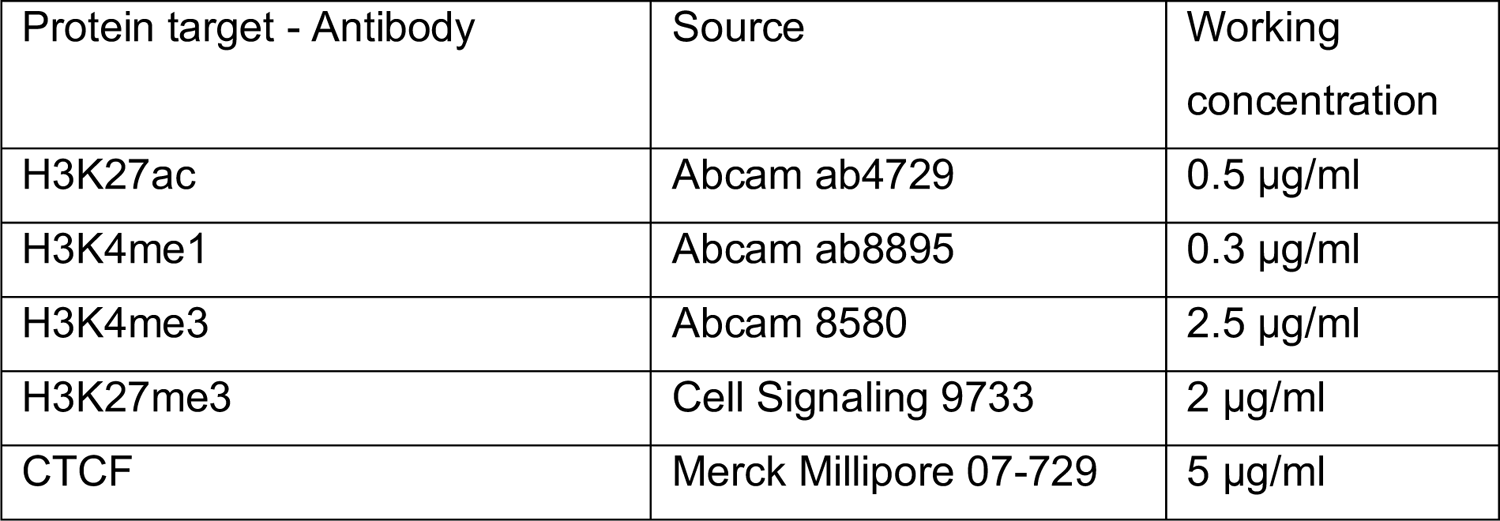
Antibodies used for ChIP-seq

| Protein target - Antibody | Source | Working concentration |
| --- | --- | --- |
| H3K27ac | Abcam ab4729 | 0.5 µg/ml |
| H3K4me1 | Abcam ab8895 | 0.3 µg/ml |
| H3K4me3 | Abcam 8580 | 2.5 µg/ml |
| H3K27me3 | Cell Signaling 9733 | 2 µg/ml |
| CTCF | Merck Millipore 07-729 | 5 µg/ml |

**Table 6.**
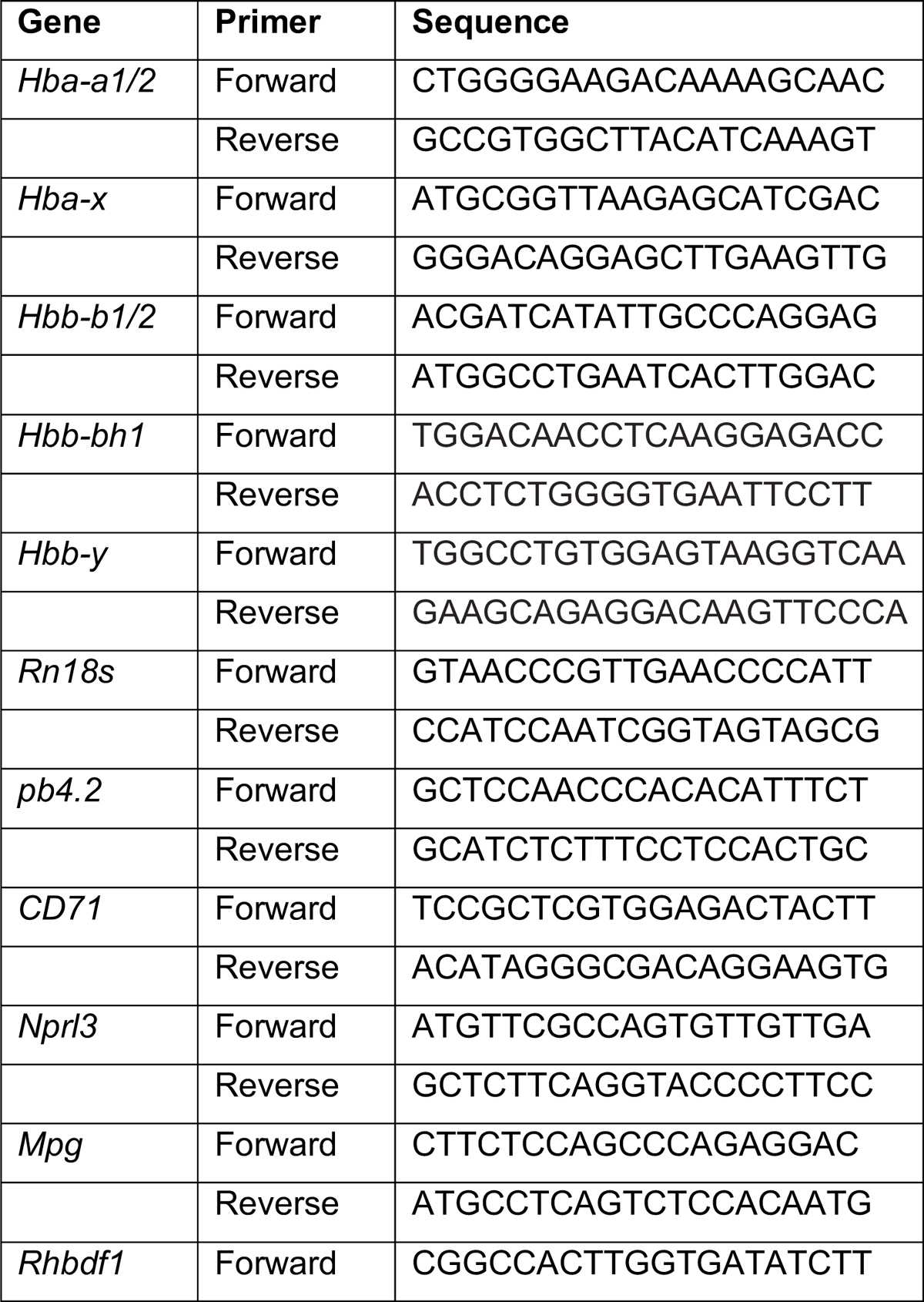
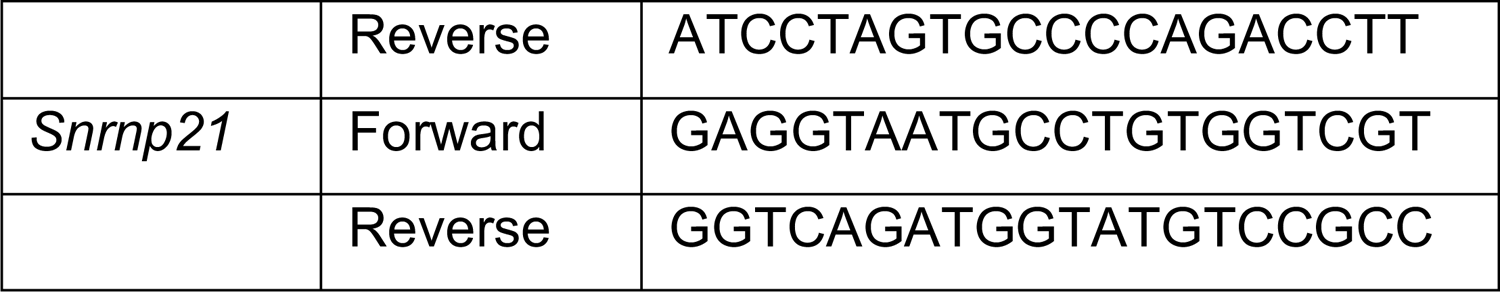
Primer pairs used for expression analysis by RT-PCR.

---

**Methods Figure 1.**
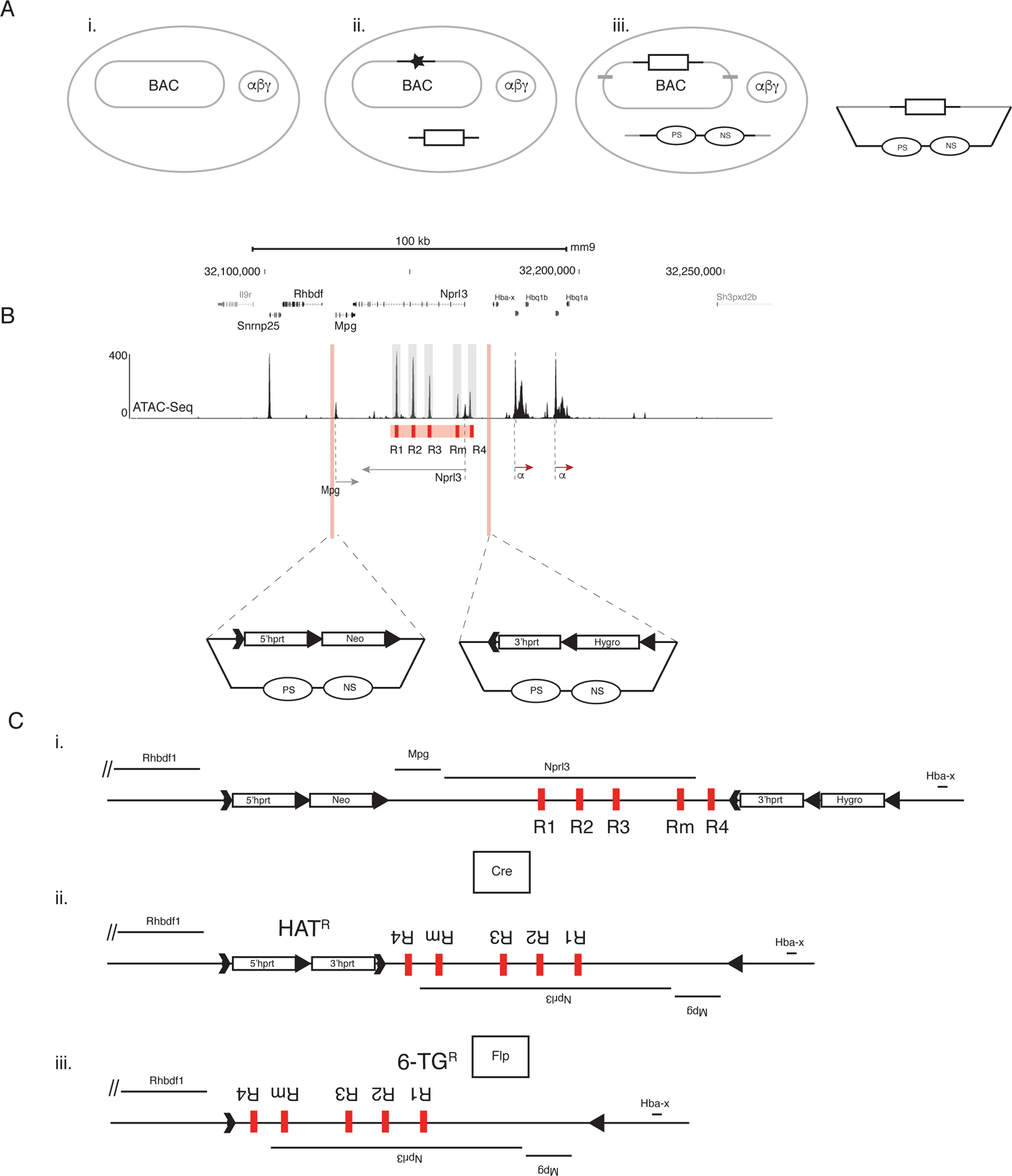
Overview of the enhancer inversion strategy. (A) Targeting vector construction using recombineering methodology. i) E.Coli containing BAC with the sequences to be targeted for LoxP insertion (upstream and downstream of Mpg and Nprl3 genes respectively) and phage-derived proteins (αβγ) necessary for target DNA alteration by homologous recombination. ii) Insertion of the LoxP sequences in the designated location in the locus (black star) using a selection cassette flanked by homology sequences to the insertion site (white box with 2 black lines). iii) Targeting vector assembly by retrieving the modified sequence using a minimal vector with positive (PS) and negative selection (NS) markers and homology sequences (grey lines) to the boundary of the homology arms required to target the locus in ES cells. (B) LoxP insertion at two positions in the α-globin locus in mouse ES cells. Schematic representation of the mouse α-globin locus and upstream region (genes and enhancers annotated as in Fig1-b). Two targeting vectors harbouring arms of homology (3 to 5 Kb) to sequences flanking the LoxP insertion sites. Selection cassettes flanked by LoxP and Frt sites in opposite directions are used. (C) Cre-mediated inversion of the region encompassing the enhancers. i) Schematic representation of a targeted allele in cis, ii) the inversion of the locus and deletion of positive selection cassettes upon Cre-recombinase expression (selected for by the restoration of the hprt gene expression, HATR), iii) Flp-mediated deletion of hprt gene (selected for by resistance to 6-TG upon HAT deletion, 6-TGR).

**Methods Figure 2.**
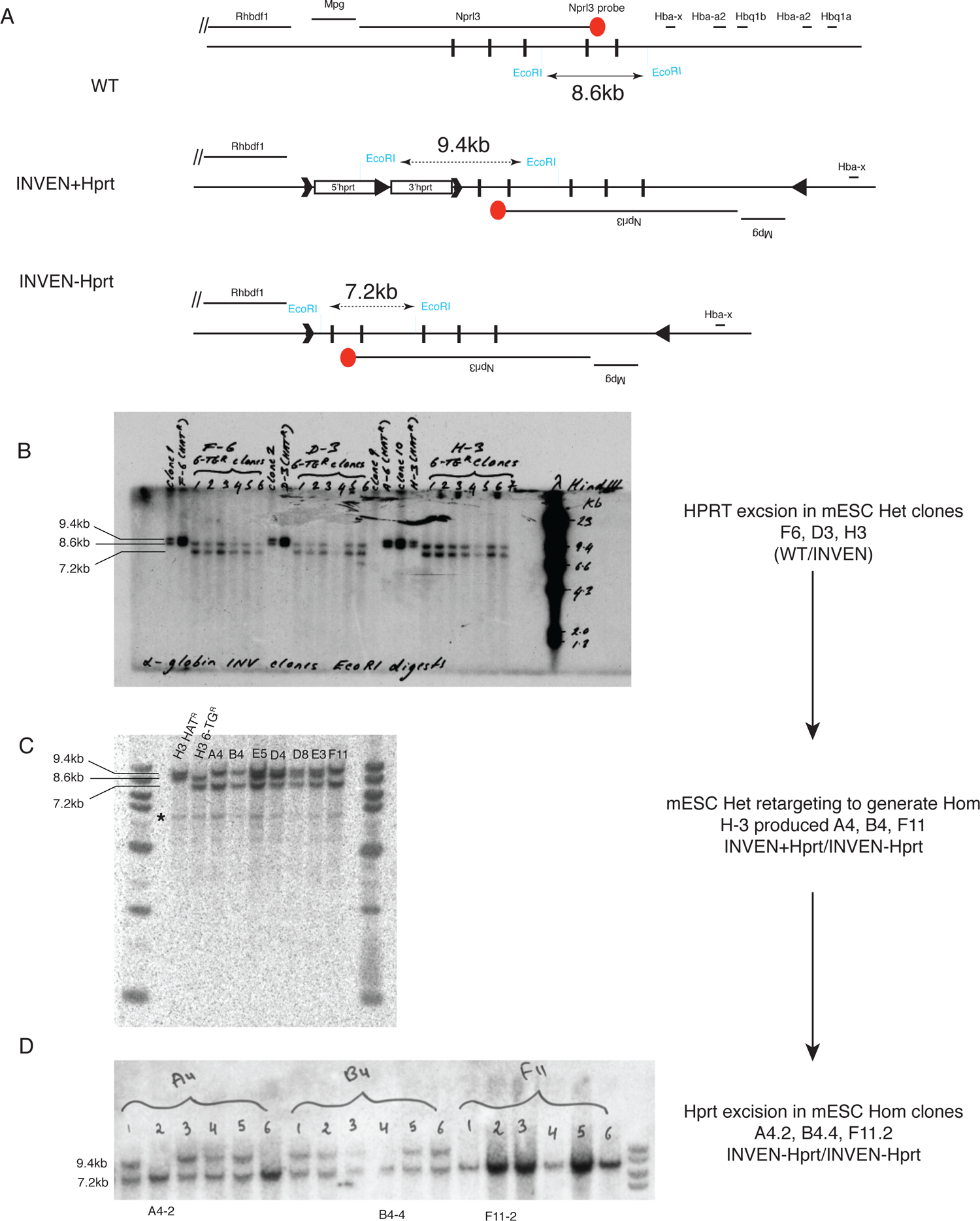
Southern blot data showing heterozygous (WT/INVEN) and homozygous INVEN (INVEN/INVEN) mESCs clones before and after Hprt minigene excision. (A) Schematic representation of the Southern Blot screening strategy for Hprt minigene excision upon flp expression. The horizontal black line indicates the genomic region spanning the α-globin locus with genes annotated on top and marked by smaller horizontal black lines. The red circle indicates the location of the probe designed within the Nprl3 gene. Vertical boxes indicate the 5 enhancers (R1, R2, R3, Rm, R4 from 3’ to 5’). EcoRI restriction enzyme digestion fragment is designated by a double-arrow line (8.6kb fragment in WT, 9.4Kb in INVEN allele with Hprt minigene intact (INVEN+Hprt), 7.2Kb in INVEN allele with Hprt minigene excised (INVEN-Hprt)). (B) Southern blot confirming Hprt minigene excision in 3 independent mESCs heterozygous for INVEN (WT/INVEN); F6, D3, H3. Note the change in the size of the bands from 9.4kb (unexcised) to 7.2Kb (excised) in 6 subclones for each of the three clones. Lambda phage DNA digested with HindIII determined the large genomic fragment sizes (ladder to the right of the blot). (C) Southern blot confirming the generation of homozygous mESC clones from a heterozygous WT/INVEN clone, already Hprt excised (H3-6TG^R^). The resulting homozygous mESCs (example A4, B4, F11) harbour one INVEN allele with excised Hprt and one INVEN allele that still retains the Hprt allele (INVEN+Hprt/INVEN-Hprt) as indicated by the size of the detected bands (9.4Kb and 7.2Kb respectively). (D) Southern blot confirming Hprt minigene excision in 3 independent mESCs (A4.2, B4.4, F11.2) homozygous for INVEN (INVEN/INVEN).

